# Inference and visualization of complex genotype-phenotype maps with *gpmap-tools*

**DOI:** 10.1101/2025.03.09.642267

**Authors:** Carlos Martí-Gómez, Juannan Zhou, Wei-Chia Chen, Arlin Stoltzfus, Justin B. Kinney, David M. McCandlish

## Abstract

Understanding how biological sequences give rise to observable traits, that is, how genotype maps to phenotype, is a central goal in biology. Yet our knowledge of genotype-phenotype maps in natural systems is limited due to the high dimensionality of sequence space and the context-dependent effects of mutations. The emergence of Multiplex assays of variant effect (MAVEs), along with large collections of natural sequences, offer new opportunities to empirically characterize these maps at an unprecedented scale. However, tools for statistical and exploratory analysis of these high-dimensional data are still needed. To address this gap, we developed *gpmaptools* https://github.com/cmarti/gpmap-tools), a *python* library that integrates a series of models for inference, phenotypic imputation, and error estimation from MAVE data or collections of natural sequences in the presence of genetic interactions of every possible order. *gpmap-tools* also provides methods for summarizing patterns of epistasis and visualization of genotype-phenotype maps containing up to millions of genotypes. To demonstrate its utility, we used *gpmap-tools* to infer genotype-phenotype maps containing 262,144 variants of the Shine-Dalgarno sequence from both genomic 5’UTR sequences and experimental MAVE data. Visualization of the inferred landscapes consistently revealed high-fitness ridges that link core motifs at different distances from the start codon. In summary, *gpmap-tools* provides a flexible, interpretable framework for studying complex genotype-phenotype maps, opening new avenues for understanding the architecture of genetic interactions and their evolutionary consequences.

## Introduction

The genotype-phenotype map describes how changes in biological sequences, such as DNA, RNA or proteins, give rise to variation in observable traits. Understanding this relationship is essential across many areas of biology, ranging from evolutionary theory (Wright, 1932; Kondrashov *et al*., 2002; Phillips, 2008; Weinreich *et al*., 2013; De Visser and Krug, 2014; Sailer and Harms, 2017; Bank, 2022; Johnson *et al*., 2023) and human disease (Moore and Williams, 2009; Dasari *et al*., 2021; Moulana *et al*., 2023a), to synthetic biology, protein engineering applications (Yang *et al*., 2019; Freschlin *et al*., 2022; Lipsh-Sokolik and Fleishman, 2024) and plant and animal breeding (De Los Campos *et al*., 2013; Sackton and Hartl, 2016; Soyk *et al*., 2020; Dwivedi *et al*., 2024).

However, our current understanding of genotype-phenotype maps in nature remains limited due to two fundamental challenges. First, the number of possible genotypes is astronomically large, making it impossible to explore more than a tiny fraction of this space in any empirical setting. Second, the phenotypic effect of a mutation often depends on the genetic background in which it occurs, a phenomenon known as epistasis (Starr and Thornton, 2016; Domingo *et al*., 2019; Miton *et al*., 2021; Bank, 2022). Addressing this context dependence requires experimental and computational approaches that are capable of capturing these complex genetic interactions.

One powerful approach to study empirical genotypephenotype maps is to experimentally construct sequence variants and measure their functional consequences. Historically, these studies were limited by the difficulty of engineering a large number of genetic variants and quantifying their phenotypes at scale, restricting most empirical genotype-phenotype maps to small numbers of genotypes, typically ranging from tens to a few hundreds (Khan *et al*., 2011; Chou *et al*., 2011; Flynn *et al*., 2013; Szendro *et al*., 2013; Ogbunugafor *et al*., 2016; Weinreich *et al*., 2018; Gao *et al*., 2022; Aguirre *et al*., 2023; Zebell *et al*., 2025). The development of Multiplex Assays of Variant Effects (MAVEs) (Kinney *et al*., 2010; Fowler and Fields, 2014; Kinney and McCandlish, 2019) has increased our phenotyping throughput by several orders of magnitude, enabling the simultaneous measurement of libraries containing thousands to millions of genotypes in a single experiment. These techniques have been used to characterize the phenotypic landscapes for short regulatory elements (Noderer *et al*., 2014; Bonde *et al*., 2016; Wong *et al*., 2018; Kuo *et al*., 2020; Komarova *et al*., 2020; Westmann *et al*., 2024b,a; Kuo *et al*., 2025; Chattopadhyay *et al*., 2025), RNAs (Domingo *et al*., 2018; Baeza-Centurion *et al*., 2019; Bendixsen *et al*., 2019; Soo *et al*., 2021; Rotrattanadumrong and Yokobayashi, 2022) and proteins (O’Maille *et al*., 2008; Bank *et al*., 2016; Wu *et al*., 2016; Starr *et al*., 2017; Poelwijk *et al*., 2019; Lite *et al*., 2020; Jalal *et al*., 2020; Moulana *et al*., 2023b; Papkou *et al*., 2023; Sundar *et al*., 2024; Zarin and Lehner, 2024; Escobedo *et al*., 2024; Johnston *et al*., 2024; Herrera-Álvarez *et al*., 2025), and combinatorial gene interactions (Bakerlee *et al*., 2022). Yet the highly combinatorial nature of these data poses significant challenges, and accurate inference typically relies on complex latent-variable models (Bloom, 2015; Otwinowski *et al*., 2018; Tareen *et al*., 2020; Tonner *et al*., 2022; Faure and Lehner, 2024) or neural networks (Bryant *et al*., 2021; Gelman *et al*., 2021; Sethi and Zhou, 2024). Gaussian processes offer an alternative class of flexible models that both capture high order interactions and provide accurate uncertainty quantification (Romero *et al*., 2013; Yang *et al*., 2019; Zhou and McCandlish, 2020; Zhou *et al*., 2022, 2025; Petti *et al*., 2025). Moreover, the mathematical tractability of Gaussian processes means that they can be designed and interpreted in terms of existing genetic concepts, for example by explicitly learning the variance explained by epistatic interactions of different orders, which was shown to allow accurate inference and statistical analysis of complex genotype-phenotype maps from MAVE data (Zhou *et al*., 2022).

An alternative approach to study genotype-phenotype maps consists in analyzing collections of natural sequences. Since natural selection tends to preserve functional sequences, we can assume that the probability of observing a given sequence in nature depends on how well it performs its function. Thus, the probability distribution over sequences with a shared function can be interpreted as a genotypephenotype map, where the phenotype is the probability of observing a sequence. Independent site models, such as Position-Weight Matrices (PWMs), estimate this sequence probability distribution by assuming that positions are independent from each other (Stormo, 2013), whereas pairwise interaction models, also known as Potts models, relax this assumption by allowing interactions between pairs of positions (Sly, 2011; Morcos *et al*., 2011; Ekeberg *et al*., 2013; Stein *et al*., 2015; Haldane and Levy, 2021). These models have proven very effective in predicting structural contacts in proteins (Marks *et al*., 2012; Haldane *et al*., 2016, 2018) and between proteins (Bitbol *et al*., 2016; Malinverni and Babu, 2023), as well as for predicting the effects of mutations in human proteins (Hopf *et al*., 2017). Pairwise interaction models have also been useful for identifying novel functional proteins (Russ *et al*., 2020) and regulatory sequences (Yeo and Burge, 2004), and for quantifying the strength of selection at the gene level (Vigué and Tenaillon, 2023). A recently proposed Gaussian process model further generalizes these methods by inferring sequence probability distributions under a prior that controls the magnitude of local epistatic coefficients (Chen *et al*., 2021, 2024). These Gaussian process based generalizations of pairwise interaction models open the opportunity to study complex genotype-phenotype maps containing higher-order epistatic interactions using readily available collections of natural sequences.

Another important challenge is the interpretation of complex genotype-phenotype maps. One way to develop an intuitive understanding of complex datasets is through data visualization tools. An approach to visualizing empirical genotype-phenotype maps is to embed the Hamming graph representing it, where each node is a genotype and edges represent single-point mutations, into a low-dimensional space. For instance, one can embed the graph by placing genotypes according to their Hamming distance to a reference sequence on one axis and their phenotype on the other (Wright, 1932; Brouillet *et al*., 2015; Domingo *et al*., 2018; Bendixsen *et al*., 2019; Baeza-Centurion *et al*., 2019; Escobedo *et al*., 2024), or applying spectral and force-directed layouts (Starr *et al*., 2017; Fragata *et al*., 2019; Martin and Ahnert, 2022; Herrera-Álvarez *et al*., 2025). However, how these representations relate to the conceptual framework of fitness peaks, valleys, and plateaus, which has shaped much of our theoretical understanding of genotype-phenotype maps (Wright, 1932; de Visser *et al*., 2018), remains unclear. A different strategy is to construct a low-dimensional representation that reflects the evolutionary dynamics induced by the genotype-phenotype map of interest. This can be done, for example, by having the distances between genotypes represent the expected time to evolve between them for a population under selection for high phenotypic values (McCandlish, 2011). This property is very useful because it places sets of functional sequences that are inaccessible to each other i.e. peaks, far apart in the visualization, naturally displaying the key genetic interactions separating them, i.e. valleys. This technique has been successfully applied to uncover qualitative features of multiple genotype-phenotype maps (Zhou and McCandlish, 2020; Zhou *et al*., 2022; Chen *et al*., 2021; Weinstein *et al*., 2023; Avizemer *et al*., 2025), as well as to study how the structure of the genetic code influences protein evolution (Rozhoňová *et al*., 2024), illustrating its potential as a general framework for interpreting and comparing complex genotype-phenotype relationships.

Here, we present *gpmap-tools*, a *python* library that offers an integrated interface to methods for inference, statistical analysis and visualization of large, complex genotypephenotype maps (Figure 1). Among its new features and other improvements, *gpmap-tools* is built on a new computational back-end that represents large matrices as linear operators, enabling memory-efficient computation. This design also allows users to simulate large genotype-phenotype maps with different types and magnitudes of epistasis, and to perform statistical analysis of quantities of interest, such as mutational effects and epistatic coefficients, even in the presence of missing data and experimental noise. *gpmap-tools* also enables calculation of the variance explained by genetic interactions involving specific sets of sites, providing a powerful tool to characterize the structure and complexity of genetic interactions. Finally, *gpmap-tools* provides an extended interface for visualizing genotype-phenotype maps with varying numbers of alleles per site, new tools for investigating the sequence features that distinguish different regions of the representation, and accelerated rendering of plots containing millions of genotypes, enabling exploration of complex genotype-phenotype maps at unprecedented scale and resolution.

**Fig. 1.**
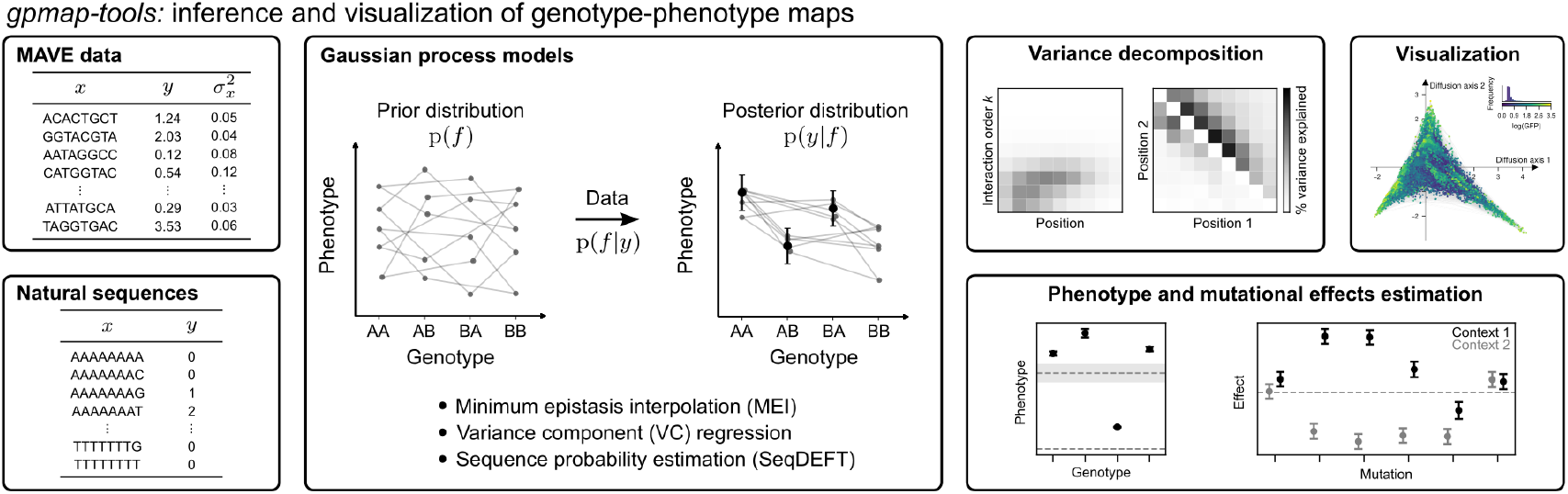
Overview of the functionality provided by *gpmap-tools*. The software uses data from Multiplex Assays of Variant Effects (MAVEs) or natural sequence variation to fit Gaussian process models for inference of empirical genotype-phenotype maps. It enables interpretation of the inferred maps by decomposing phenotypic variance into components associated with interactions of different orders and specific subsets of sites, visualizing the full genotype-phenotype landscape, and computing posterior distributions for specific genotypes or mutational effects of interest.

We demonstrate the capabilities of *gpmap-tools* by inferring the fitness landscape of the Shine-Dalgarno (SD) sequence from two fundamentally different types of data: (i) natural sequence diversity in genomic 5’ untranslated regions (UTRs) and (ii) MAVE data (Kuo *et al*., 2020). The inferred landscapes reveal a shared structure consisting of peaks corresponding to 16S rRNA binding at different distances rel-ative to the start codon. These peaks are connected by extended ridges of functional sequences when the corresponding binding sites are spaced three nucleotides apart. These ridges arise from overlapping SD motifs in the same sequence, a consequence of the motif’s quasi-repetitive structure, which allows new binding sites to emerge at offset positions without disrupting functionality. Building on this qualitative understanding of the genotype-phenotype map, we fit a simplified mechanistic model with parameters that have clear biophysical interpretations. This model allows us to disentangle the effects of mutations on binding at different registers relative to the start codon *in vivo*, while capturing the key structural features of the empirical landscape. Taken together, our analysis illustrates how *gpmap-tools* enables the inference and characterization of genotype-phenotype maps from diverse data sources, facilitating the discovery of simple molecular mechanisms capable of generating the observed architecture of epistatic interactions and offering insights into the evolutionary consequences induced by these complex genotype-phenotype maps.

### Approach

In this section, we provide a brief technical overview of the methods for inference and interpretation of genotypephenotype maps implemented in *gpmap-tools* (Figure 1), for application see Results.

A genotype-phenotype map is a function that assigns a phenotype, typically a scalar value, to every possible sequence of length ℓ on *α* alleles (where e.g. *α* =4 for DNA and *α* = 20 for proteins). This function can be represented by an *α*^ℓ^-dimensional vector *f* containing the phenotype for every possible genotype. We begin by discussing several methods for quantifying the amount and type of epistasis present in any particular vector *f*.

#### Epistasis in genotype-phenotype maps

*gpmap-tools* implements two different methods for measuring the amount and pattern of epistasis in a given genotype-phenotype map: one based on the typical magnitude of local epistatic coefficients across all possible subsets of mutations and the other based on the proportion of phenotypic variance explained by genetic interactions of different orders or involving specific subsets of sites.

##### Local epistatic coefficients

The traditional epistatic coefficient quantifies how much the effect of a mutation *A*→ *a* in one site changes in the presence of an additional mutation *B*→ *b* in a second site in an otherwise identical genetic background *C*:

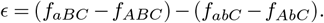

The average squared epistatic coefficient 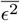 across all possible pairs of mutations and genetic backgrounds provides a measure of the variability in mutational effects between neighboring genotypes across the whole genotype-phenotype map (Zhou and McCandlish, 2020). 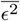 can be efficiently calculated as a quadratic form 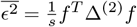, where *s* is the number of epistatic coefficients and Δ^(2)^ is a previously described sparse *α* ^*ℓ*^ ×*α* ^*ℓ*^ positive semi-definite matrix (Zhou and McCandlish, 2020). This statistic can be generalized to characterize the typical size of local *P* -way epistatic interactions (Chen *et al*., 2021) and to a setting in which alleles are naturally ordered e.g. copy number (Chen *et al*., 2024).

##### Variance components

A genotype-phenotype map *f* can be decomposed into the contributions of 𝓁 +1 orthogonal subspaces *f* = ∑_*k*_ *f*_*k*_, where *f*_*k*_ represents a function containing epistatic interactions solely of order *k*. These orthogonal components *f*_*k*_ are obtained by projecting *f* onto the th order subspace using the projection matrix *P*_*k*_. This decomposition enables quantification of the variance explained by interactions of different orders, providing a global summary of the complexity of genetic interactions in a genotypephenotype map (Stadler, 1996; Happel and Stadler, 1996; Stadler and Happel, 1999; Stadler, 2002; Zhou *et al*., 2022).

Here we show that each *f*_*k*_ can be further decomposed into the contribution of 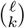 smaller orthogonal subspaces *f*_*k=*_∑_*U* :|*U* |=*k*_ *f*_*U*_, where *f*_*U*_ represents a function containing genetic interactions only among the *k* sites in *U*. These orthogonal components *f*_*U*_ are obtained by projecting the function *f* into the corresponding subspace using the orthogonal projection matrix *P*_*U*_ given by:

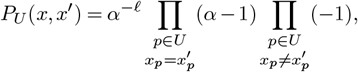

for any pair of sequences *x, x* ^′^ (see Supplementary Information A). A similar decomposition was shown for certain models of sequence-function relationships parametrized using a specific scheme of interaction terms (Park *et al*., 2024; Posfai *et al*., 2025), but this formulation allows the direct decomposition of the function *f* rather than relying on a particular parameterization.

These projection matrices allow us to quantify not only the variance explained by interactions of different order, but also the variance explained by all possible subsets of sites *U*. Although there are 2^𝓁^ such subsets in total, they can be aggregated to yield informative, low-dimensional summary statistics. For instance, we can compute the variance explained by order-*k* epistatic interactions involving a specific site or pair of sites, or the variance explained by interactions of all orders involving each pair of sites (see Supplementary Information B) (Crawford *et al*., 2017; Reddy and Desai, 2021). Through these capabilities, *gpmap-tools* enables a fine-grained decomposition of epistatic variance, offering new insights into the structure and complexity of genetic interactions across sites.

#### Gaussian process inference of genotype-phenotype maps

Gaussian process models are a class of Bayesian non-parametric models that place a multivariate Gaussian prior distribution over all possible functions and compute the posterior distribution given observed data (Rasmussen and Williams, 2008). In our case, we assign a zero-mean Gaussian prior distribution over genotype-phenotype maps p(*f*) characterized by either its covariance matrix *K* or precision matrix *C*. The covariance matrix *K* is most often defined through a kernel function that returns the prior covariance between any pair of sequences. Then, given some data *y* and using a likelihood function p(*y* |*f*), we update the probability distribution of plausible genotype-phenotype maps to be consistent with these observations by computing the posterior distribution p(*f* |*y*).

##### Interpretable priors

*gpmap-tools* implements two families of priors based on the two approaches to quantify epistasis in genotype-phenotype maps described above: one family that is defined in terms of local epistatic coefficients and a second that is defined in terms of variance components. The first prior is parametrized by its precision matrix 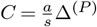 and assigns a prior probability to *f* depending on its average squared epistatic coefficient of order 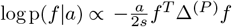 (Zhou and McCandlish, 2020; Chen *et al*., 2021). This prior implicitly leaves genetic interactions of order *k*< *P* unconstrained, and hence correspond to the use of an improper Gaussian prior. For examples, for *P* = 2 additive effects are not penalized, for *P* =3 additive and pairwise effects are not penalized, etc. For fixed *P*, this family of priors has a single hyperparameter *a* that is inversely proportional to the expected average squared local epistatic coefficient under the prior. As *a*→ 0, we assign the same prior probability to every possible genotype-phenotype map. On the other hand, as *a*→∞, we decrease the prior probability of genotype-phenotype maps with non-zero local *P* -epistatic coefficients (Chen *et al*., 2021).

The second family of priors are the variance component priors, which are parametrized by their covariance matrix 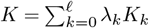, where *K*_*k*_ represents the covariance matrix for genotype-phenotype maps with only *k*th order interactions. The ℓ +1 hyperparameters *λ*_*k*_ control the variance explained by genetic interactions of order *k* (Neidhart *et al*., 2013) and equivalently the decay in the predictability of mutational effects and epistatic coefficients as the number of mutations separating two genetic backgrounds increases (Zhou *et al*., 2022). The formal relationship between the two sets of priors is that the priors based on the Δ^(*P*)^ operators can be obtained as limits of the variance component prior (Zhou *et al*., 2022).

These prior distributions for *f* have hyperparameters with clear biological interpretations in terms of the expected magnitude and type of epistasis. This allows users to define the corresponding priors in a principled and interpretable way. Moreover, under the assumption that the structure of epistasis observed in the data generalizes to the full genotypephenotype map, *gpmap-tools* can infer these hyperparameters using either cross-validation or kernel alignment (Rasmussen and Williams, 2008; Wang *et al*., 2015), providing estimates that are both data-driven and biologically meaningful.

##### Likelihood functions

*gpmap-tools* implements two likelihood functions for inference of genotype-phenotype maps from different types of data. For experimental data, the vector *y* contains the measurements associated to a subset of sequences *x* with known Gaussian measurement variance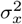.

Thus, the likelihood function is given by

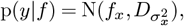

where 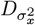 is a diagonal matrix with 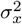 along the diagonal. For observations of natural sequences, data consists of the number of times *N*_*i*_ a given sequence *i* was observed out of a total of *N*_*T*_ = _*i*_ *N*_*i*_ observations. In this case, the likelihood function is given by the multinomial distribution

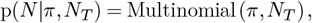

where *π* is the vector representing the sequence probability distribution, such that *π*_*i*_ is corresponds to the probability of observing sequence *i*.

##### Posterior distributions

*gpmap-tools* enables the computation of the posterior distribution over the space of possible genotype-phenotype maps using both Gaussian and multinomial likelihood functions. Under a Gaussian likelihood, the posterior distribution is also a Gaussian 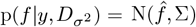 with closed form analytical solutions for the mean 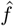 and covariance matrix ∑. *gpmap-tools* implements the classical solution expressed in terms of the prior covariance matrix *K* (Rasmussen and Williams, 2008) but also the solution when the prior is defined by its precision matrix *C*, which are equivalent when *C* = *K*^−1^ (see Supplementary Information F).

Under non-Gaussian likelihood functions, such as the multinomial likelihood, the posterior distribution has no closed form analytical solution. However, given that log p(f |y) is proportional to log p(y| f) + log p(f), which can be efficiently computed for any *f*, *gpmap-tools* leverages optimization methods to find the maximum a posteriori (MAP) 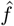 and compute an approximate Gaussian posterior using the Laplace approximation, where the posterior covariance matrix ∑ is defined by the inverse Hessian of the posterior at its mean 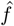 (Rasmussen and Williams, 2008).

These solutions are completely general, in the sense that they hold for arbitrary valid prior covariance of precision matrices. However, as we will explain below, *gpmap-tools* implements highly optimized versions of these calculations that take advantage of the structure of sequence space and our specific choices for *C* and *K*.

#### Inference of genotype-phenotype maps with *gpmap– tools*

*gpmap-tools* combines prior distributions with likelihood functions into a number of Gaussian process models for inference of complete genotype-phenotype maps.

##### Minimum epistasis interpolation

The minimum epistasis interpolation (MEI) method was originally proposed in terms of finding the *f*_*z*_ at unobserved sequences *z* given the known phenotype *f*_*x*_ at sequences *x* by minimizing the average squared epistatic coefficient 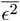 over the complete genotypephenotype map (Zhou and McCandlish, 2020). *gpmap-tools* provides a generalization to local epistatic coefficients of any order *P*, (by minimizing *f* ^*T*^ Δ^(*P*)^*f* (Chen *et al*., 2021)), incorporates known Gaussian measurement noise through 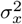 and, by re-framing minimum epistasis interpolation as a Gaussian process model, enables uncertainty quantification via the posterior covariance (see Supplementary Information E,F).

##### Empirical variance component regression

Empirical variance component regression (VC regression), proposed in (Zhou *et al*., 2022), combines a variance component prior parameterized by the variance *λ*_*k*_ associated to interactions of each possible order *k* with a Gaussian likelihood with known noise variance 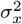 to compute the exact Gaussian posterior distribution over *f*. The hyperparameters *λ*_*k*_ controlling the expected variance explained by interactions of order *k* under the prior are optimized through kernel alignment (Wang *et al*., 2015). This procedure minimizes the squared distance between the covariance under the prior and the empirical distance-covariance function computed from the incomplete data. While a naive kernel alignment implementation requires computation with large covariance matrices, the prior covariance between two sequences depends only on the Hamming distance between them, resulting in only *ℓ* +1 different values. As a consequence, the problem of kernel alignment can be efficiently solved as a simpler *ℓ* + 1-dimensional constrained weighted least squares problem (Zhou *et al*., 2022).

##### Sequence probability distribution estimation

*gpmap-tools* also implements the SeqDEFT method for estimating probability distributions *π* over sequence space (Chen *et al*., 2021).

SeqDEFT parametrizes the probability distribution as

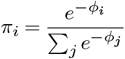

and defines an improper prior distribution over the latent phenotype *φ* that penalizes local epistatic coefficients of order *P* given by log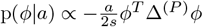, which is combined with a multinomial likelihood function to compute an approximate posterior distribution over *φ* = −log *π*. The hyperparameter *a* is optimized by maximizing the crossvalidated log-likelihood under the MAP estimate over a onedimensional grid search. This also enables to examination of the behavior of the model towards the two limiting solutions i.e. when local epistatic coefficients are unconstrained (*a* = 0) or forced to be zero (*a* → ∞) (Chen *et al*., 2021).

#### Visualization of genotype-phenotype maps

Genotypephenotype maps are inherently high-dimensional objects, and thus difficult to visualize in an intuitive manner. *gpmap-tools* implements a previously proposed strategy for visualizing fitness landscapes (McCandlish, 2011) that computes embedding coordinates for genotypes such that squared distances between pairs of genotypes in the low-dimensional representation approximate the expected times to evolve from one to another under selection for high phenotypic values. This layout highlights regions of sequence space containing highly functional genotypes that are nevertheless poorly accessible to each other e.g. fitness peaks separated by valleys, or sets of sequences where the intermediates are functional but the order of the intervening mutations is highly constrained.

##### Evolutionary model

We assume a weak mutation model of evolution in haploid populations, such that mutations are always fixed or lost before a new mutation arises (McCandlish, 2011, 2018; Zhou and McCandlish, 2020; Chen *et al*., 2021). Under this model, the evolutionary rate *Q*(*i, j*) from genotype *i* to *j* depends on the mutation rate *M* (*i, j*) (which we assume is taken from a time-reversible mutational model) and the probability of fixation relative to a neutral mutation (Bulmer, 1991; McCandlish and Stoltzfus, 2014):

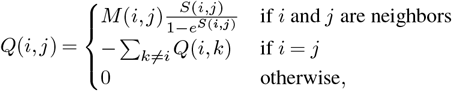

where *S*(*i, j*) is the scaled selection coefficient of genotype *j* relative to genotype *i*. For the purposes of constructing a useful visualization, we then assume that this scaled selection coefficient is proportional to the phenotypic differences between the two genotypes i.e. *S*(*i, j*)= *c*(*f* (*j*) −*f* (*i*), where the constant *c* can be interpreted as the scaled selection coefficient (2*N*_*e*_*s*, for a Haploid Wright-Fisher population) associated with a phenotypic difference of 1. Unless specifically studying the role of mutational biases on evolution on empirical landscapes, we would typically assume that *M* (*i, j*)=1 for any *i, j* pair (i.e. measuring time in units of the inverse mutation rate), and focus on the evolutionary dynamics induced by the structure of the genotype-phenotype map alone. This model assigns a low but non-zero probability of fixation to deleterious mutations and has a unique stationary distribution *π*(*i*) given by

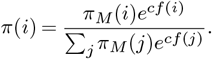

where *π*_*M*_ (*i*) are the time-reversible neutral stationary frequencies, which are uniform in absence of mutational biases (Sella and Hirsh, 2005; McCandlish *et al*., 2015). The stationary distribution can be used to select a reasonable value of *c* for our evolutionary process. When representing a probability distribution, such as one inferred using SeqDEFT, setting *f* (*i*) = log *π*(*i*) and *c* = 1 will result in a stochastic process in which the stationary distribution exactly matches the estimated genotype probabilities, providing a very natural representation of the landscape. When inferring the genotype-phenotype map from MAVE data, *c* can be adjusted so that the mean phenotype under the stationary distribution aligns with realistic natural values e.g. the phenotype associated to a wild-type or reference sequence. Alternatively, a range of *c* values can be used to generate a family of visualizations for a single genotype-phenotype map to reflect the evolutionary impact of its structure under different assumptions concerning the relative strengths of selection and drift.

##### Low-dimensional representation

The right eigenvectors *r*_*k*_ of *Q* associated to the largest eigenvalues *λ*_*k*_ (*λ*_1_ =0 > *λ*_2_ ≥*λ*_3_ ≥...) can be computed using iterative methods that leverage the sparse structure of *Q*. When appropriately normalized and re-scaled as 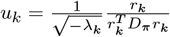, the first few *r*_*k*_ for *k*≥ 2 can be used as embedding coordinates, resulting in a low-dimensional representation in which squared distances between genotypes optimally approximate the commute times i.e. the sum of hitting times *H*(*i, j*) from *i* to *j* and *H*(*j, i*) from *j* to *i*, thus separating sets of functional genotypes that are largely inaccessible to each other for a population evolving under selection for high phenotypic values:

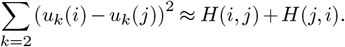

The eigenvalues *λ*_*k*_ represent the rates at which the associated eigenvectors become less relevant for predicting evolutionary outcomes with time. The associated relaxation times 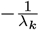 have units of expected number of substitutions and allow us to identify components that decay slower than expected under neutral evolution, where we note that if all mutations occur at rate 1, the neutral relaxation time is given by the reciprocal of the minimum number of alleles across sites. Because *u*_*k*_ captures the *k* −1-th strongest barrier to the movement of a population in sequence space, we refer to*u*_*k*_ as diffusion axis *k* −1 (see McCandlish, 2011; Chen *et al*., 2021, for more details).

##### Rendering and visualization

In addition to computing the coordinates *u*_*k*_, *gpmap-tools* provides functionality at both high and low levels to plot and render the visualizations of genotype-phenotype maps using different backend plotting libraries. This includes the standard plotting library in python, *matplotlib* (Hunter, 2007), for generating highly customized visualizations, and *plotly* (Hossain, 2019), for generating interactive 3D visualizations that display the sequence associated to each node when hovering the mouse over them. Moreover, as rendering large numbers of points and lines becomes limiting in large datasets, the *gpmap-tools* plotting library leverages the power of *datashader* (Bednar *et al*., 2022) for efficiently rendering plots containing millions of different elements, achieving close to an order of magnitude speed up for large genotype-phenotype maps (Figure S1).

#### Efficient computation with *gpmap-tools*

We aim to study genotype-phenotype maps with a number of genotypes ranging from a few thousands up to millions. However, all of the described methods require computing with unreasonably large matrices of size *α*^*ℓ*^× *α*^*ℓ*^. For instance, to study a genotype-phenotype map for 9 nucleotides, a naive implementation would need to build a 4^9^ ×4^9^ matrix requiring 512GB of memory using 64 bit floating point numbers and over 100 billion operations to compute matrix-vector products. While some of the necessary matrices are sparse e.g. Δ^(*P*)^ and *Q*, allowing efficient storage and computation (McCandlish, 2011; Zhou and McCandlish, 2020; Chen *et al*., 2021), other matrices e.g. *P*_*U*_ and *K*, are dense.

*gpmap-tools* circumvents these challenges using two strategies. First, we note that every matrix *A* with entries *A*_*ij*_ depending only on the Hamming distance between sequence *i* and *j*, such as Δ^(*P*)^ as well as the dense matrices *P*_*k*_ and *K*_*k*_, can be expressed as an *ℓ*-order polynomial in the Laplacian of the Hamming graph *L* (Zhou *et al*., 2022). This enables efficient computation of matrix-vector products 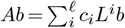 by multiplying the vector *b* by *L* up to *ℓ* times, e.g. *L*^2^*b*= *L*(*Lb*), and taking linear combinations of the results without explicitly building the possibly dense matrix *A. gpmap-tools* also implements *L* as a linear operator (see Supplementary Information D). While this new linear operator-based implementation achieves comparable efficiency to a sparse matrix formulation in nucleotide space, it is an order of magnitude faster in protein spaces (Figure S2). More importantly, the *L* linear operator requires virtually no time for construction and has much lower memory requirements (Figure S2).

Second, we note that many of the relevant matrices can be obtained as *ℓ*-Kronecker products of *α* × *α* matrices, such as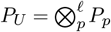. By using *scipy*’s (Virtanen *et al*., 2020) LinearOperators functionality, we can leverage the mixed Kro-necker matrix-vector product property to enable computation of e.g. *P*_*U*_ *b* without constructing *P*_*U*_ (see Supplementary Information C, Figure S3). Rather than calculating explicit inverse matrices, we can likewise use these linear operators to find numerical solutions to matrix equations using Conjugate Gradient (CG). By combining multiple linear operators, we are able to compute the posterior variance for a small number of sequences of interest or the posterior covariance for any set of linear combinations of phenotypic outcomes e.g. calculating posterior variance for mutational effects in specific genetic backgrounds and epistatic coefficients of any order, while limiting the number of linear systems to solve with CG to the number of linear combinations of interest.

## Results

In this section, we illustrate the power of *gpmap-tools* by studying the genotype-phenotype map of the Shine-Dalgarno (SD) sequence. The SD sequence is a motif located in the 5’UTR of most prokaryotic mRNAs. This motif is recognized by the 3’ tail of the 16S rRNA through base pair complementarity with a region known as the anti Shine-Dalgarno (aSD) sequence, promoting translation initiation (Shine and Dalgarno, 1975). Understanding how the SD sequence modulates protein translation *in vivo* is key in synthetic biology applications (Salis *et al*., 2009; Gilliot and Gorochowski, 2024). Previous studies used existing sequence diversity (Hockenberry *et al*., 2018; Wen *et al*., 2020) and MAVE data (Bonde *et al*., 2016; Kuo *et al*., 2020) to build models for this genotype-phenotype map. However, these models cannot account for higher-order genetic interactions and provide limited understanding of the structure of the genotype-phenotype map. Thus, *gpmap-tools* offers a new opportunity to model and understand the patterns of genetic interactions and the main qualitative features that define this important regulatory element.

### Inferring the probability distribution of the Shine-Dalgarno sequence

Here, we use SeqDEFT to infer the sequence probability distribution for the SD sequence by using the 5’ untranslated regions (UTRs) across the whole *E. coli* genome. We extracted the 5’ UTR sequence from 5,311 annotated genes and aligned them with respect to the start codon. Figure 2A shows site-specific allele frequencies for up to 20bp upstream of the start codon.

**Fig. 2.**
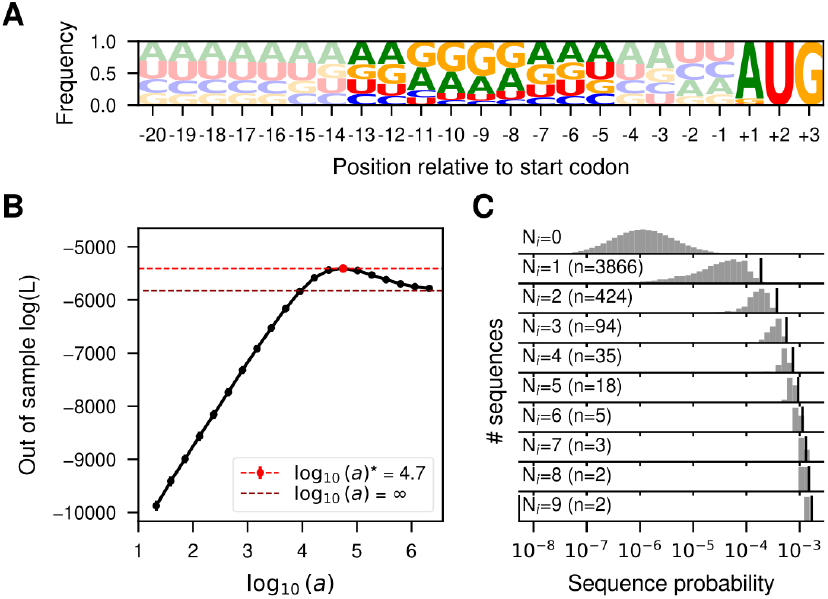
Inference of the probability distribution of the Shine-Dalgarno sequence. (A) Sequence logo representing the site-specific allele frequencies of 5,311 5’UTRs in the *E. coli* genome aligned with respect to the annotated start codon. The start codon and the 9 nucleotide region 4 bases upstream are highlighted to emphasize the region most relevant for translation initiation. (B) Log-likelihood computed in the 20% held-out sequences in 5-fold cross-validations of a series of SeqDEFT models (P=2) under varying values of the hyperparameter *a*. The horizontal dashed lines represent the log-likelihood of the limiting maximum entropy model (black), corresponding to the independent sites model shown in panel A, or the best SeqDEFT model (red). (C) Distribution of inferred sequence probabilities depending on the number of times *N*_*i*_ they were present in the *E. coli* genome represented in a logarithmic scale. Vertical black lines represent the empirical frequency *N*_*i*_*/N*_*T*_ corresponding to each *N*_*i*_ value.

These allele frequencies showed an enrichemnt in purines between positions −13 and −5 that is characteristic of the location of the SD sequence, which classically exhibits the consensus sequence AGGAGGU (Shine and Dalgarno, 1975; Hockenberry *et al*., 2018; Wen *et al*., 2020). Thus, we attempted to infer a genotype-phenotype map for this 9 nucleotide region. Out of the total 4^9^ = 262, 144 possible sequences, we observed 3, 690 unique sequences, most of them observed a single time. Given that the number of sampled sequences is two orders of magnitude smaller than the number of possible sequences, we expect many unobserved sequences to be functional. Therefore, sharing information across neighboring genotypes through SeqDEFT’s prior distribution could alleviate this limited amount of data. Figure 2B shows that the SeqDEFT model better predicts the frequencies of held-out sequences than either the site-independent model (the maximum entropy model given the site-specific frequency profiles, *a* = ∞) or as we approach the empirical frequencies model (*a* →0, which maximizes the likelihood). In particular, this optimum at an intermediate value of *a* provides strong support for the presence of epistatic interactions (Chen *et al*., 2021).

We then computed the MAP solution (using all available data) under the value *a*^*^ that maximized the likelihood for the held-out sequences and compared the inferred probabilities with the observed frequencies among *E. coli* 5’ UTRs (Figure 2C). Sequences that appear more than 2-3 times are always inferred to be highly functional (i.e. high frequency). However, there is a wide range of variability for unobserved sequences, ranging 4 orders of magnitude in their estimated probabilities, many of them with larger probabilities than some sequences that are observed once. The MAP solution yields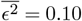, corresponding to a root mean square local epistatic coefficient of 0.32, which is slightly less than half of the size of the root mean squared mutational effect (0.78). This indicates that adding a single mutation to the genetic background often substantially changes the effects of other mutations.

### Inferring the genotype-phenotype map of the Shine– Dalgarno sequence from MAVE data

We next used data from a previously published MAVE (Kuo *et al*., 2020) measuring the expression of a GFP reporter controlled by a sequence library containing nearly all 262,144 possible 9 nucleotide sequences 4 nucleotides upstream of the start codon, i.e., the same region considered in our previous analysis. We first run MEI to predict the phenotype for all missing genotypes. The imputed genotype-phenotype map had an 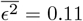. This value is not directly comparable with the results of our SeqDEFT analysis because of the difference in measurement scale (log probability vs. log GFP). Still, we can compare the root mean squared epistatic coefficient, which for MEI takes a value of 0.32, to the root mean squared size of mutational effects, which for MEI is 0.33. These similar magnitudes indicate that there is substantial variability in the effects of mutations across neighboring genotypes, more so than in the genotype-phenotype map inferred with SeqDEFT.

To better capture this high degree of inferred epistasis, we turned to VC regression, where the prior reflects the observed predictability of mutational effects in the training data. We found that the empirical phenotypic correlation between pairs of sequences decayed quite quickly with the number of mutations e.g. pairs of sequences separated by three mutations only showed a correlation of 0.25 between their measured phenotypes (Figure 3A). We next estimated the vari-ance component prior distribution that best matched the observed distance correlation patterns and computed the variance explained by interactions of every possible order under this prior (Figure 3B). The additive and pairwise component explained only 57.6% of the overall variance, suggesting an important influence of higher-order genetic interactions. We then inferred the complete genotype-phenotype map under this prior. These estimates recapitulated the experimental data remarkably well (*R*^2^ = 0.94, Figure 3C) and made predictions almost as accurate in held-out test sequences (*R*^2^ = 0.87, Figure 3D). Importantly, our estimates of the uncertainty of the phenotypic predictions are well calibrated, as we find approximately the expected fraction of measurements in the test set within posterior credible intervals (Figure S4C). Comparing the predictive performance of MEI against VC regression as a function of the number of sequences used for training, we find that while the two models perform comparably well when the genotype-phenotype map is densely sampled, and MEI performs better with extremely low sampling (likely due to error in the estimation of variance components), overall VC regression exhibited substantially higher performance across a wide range of training data densities (Figure S4A,B) and better calibration of the prediction’s uncertainty (Figure S4C).

**Fig. 3.**
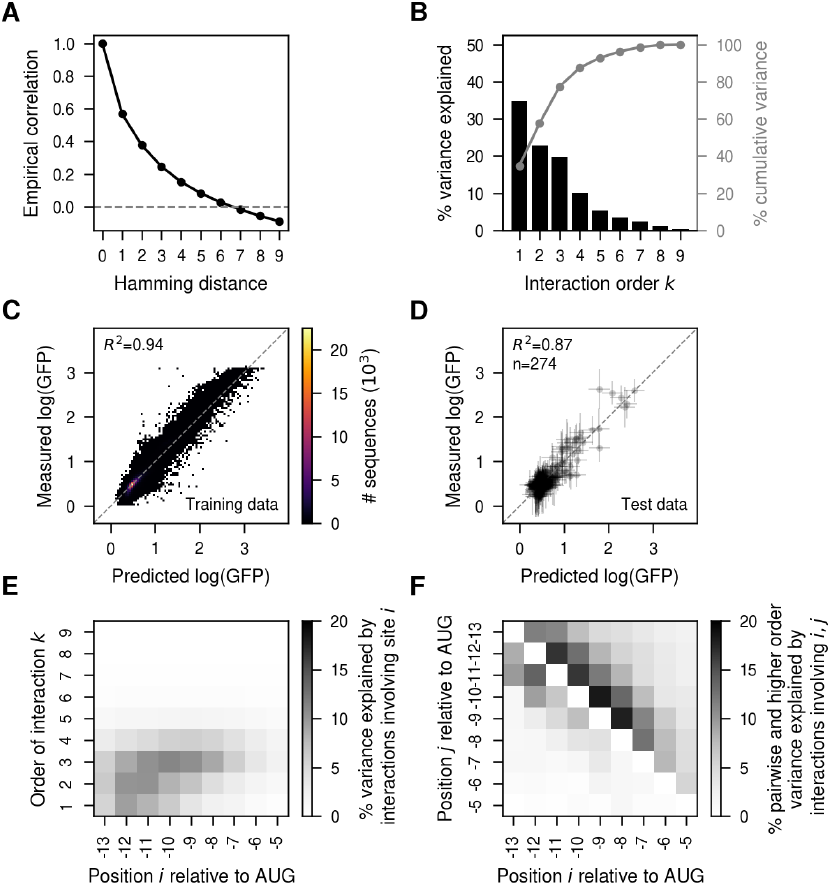
VC regression analysis of the experimentally measured genotype-phenotype map for the Shine-Dalgarno sequence in the dmsC gene context (Kuo *et al*., 2020).(A) Empirical distance-correlation function using the measured log(GFP) values in the experimentally evaluated sequences. (B) Percentage of variance explained by interactions of order *k* in the inferred VC regression prior. Grey lines represent the cumulative percentage of variance explained by interactions up to order *k*. (C) Two-dimensional histogram showing the comparison of the measured log(GFP) and the MAP estimate under the VC model in sequences used for model fitting. (D) Comparison of the posterior distribution for held-out test sequences and the measured log(GFP) values. Horizontal error bars represent posterior uncertainty represented as the 95% credible interval, whereas vertical error bars correspond to the 95% confidence interval under each measurement’s variance. (E) Heatmap representing the percentage of variance explained by interactions of order *k* involving each position relative to the start codon. (F) Heatmap representing the percentage of variance explained by pairwise (lower triangle) and higher-order (upper triangle) interactions that is explained by interactions involving pairs of positions relative to the start codon. See Supplementary Information B for details on calculating (E,F).

### Position-specific contributions to epistasis

An important advance in *gpmap-tools* is its ability to use the *P*_*U*_ matrices to evaluate the contribution of each site to genetic interactions of different orders (Supplementary Information B). Figure 3E shows this analysis for the MAP solution obtained using VC regression. We see that while positions −6 and −5 have an overall weak influence in the measured translational efficiency, and sites −13 to −10 have both strong additive and epistatic contributions, sites −9 to −7 influence the phenotype mostly through higher-order epistatic interactions. Thus, we find that sites in the SD sequence have very heterogenous contributions to genetic interactions of different orders, with some sites having stronger additive and lower order epistatic interactions, whereas other sites influence translation primarily via higher-order interactions. These variances can be further decomposed into variances explained by epistatic interactions of any order involving each possible pair of sites (Supplementary Information B). This decomposition reveals that pairwise interactions are largely confined to sites within 3 nucleotides of each other and are strongest between positions −13 to −10 (Figure 3F, lower triangle). Higher-order interactions extend to sites separated by up to 4 nucleotides, with the most prominent effects involving positions −9 to −7 (Figure 3F, upper triangle). In contrast, interactions between sites separated by 5 or more nucleotides are rare across all orders (Figure 3F). These findings indicate that the effect of a mutation at a given site depends primarily on nearby sites and becomes nearly independent of mutations beyond a 4-nucleotide range. Overall, the ability to quantify the variance explained by interactions of different orders and positional combinations offers a powerful framework for characterizing the nature and strength of epistasis, and for identifying communities of interacting sites within genotype-phenotype maps.

### Visualizing the probability distribution of the SD sequence

In order to understand the main qualitative properties of this highly epistatic genotype-phenotype map, we generated a low-dimensional representation using our visualization technique. Figure 4A shows that the genotypephenotype maps consists of at least three largely isolated peaks. These peaks correspond to the canonical SD motif AGGAG located at three consecutive positions relative to the start codon, with a fourth central peak corresponding to a shift of the canonical motif one additional base upstream appearing along Diffusion axes 3 in a 3-dimensional representation (Figure S5). This shows that not only can the aSD sequence bind at different distances from the start codon to induce efficient translation initiation, consistent with the interaction neighborhoods shown in Figure 3D, but also that it is hard to evolve a sequence with a shifted SD motif by one or two positions through single point mutations without losing translational efficiency. In contrast, sequences with an SD motif shifted by three positions remain largely connected by extended ridges of functional sequences. In these extended ridges, a second binding site can evolve through a sequence of point mutations without destroying the first. Specifically, within each trinucleotide sequence around the central AGG common to two binding registers, mutations can accumulate in diverse orders, opening up many different evolutionary paths only subject to the constraint of evolving a second SD motif before destroying the first one. Figure 4A highlights two examples of such paths.

**Fig. 4.**
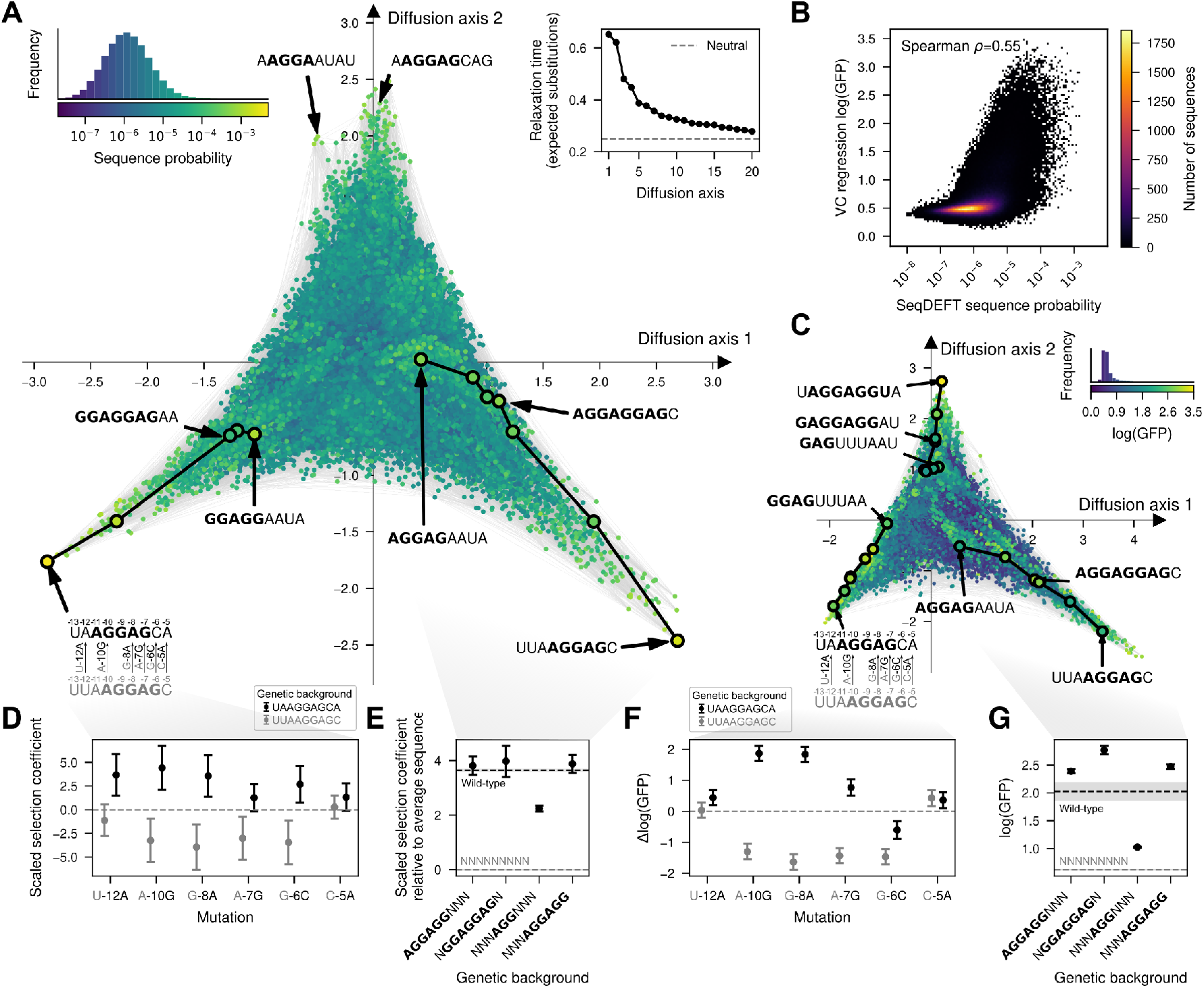
Visualization of the genotype-phenotype map of the Shine-Dalgarno sequence. (A,C) Low-dimensional representation of the *E. coli* Shine-Dalgarno sequence probability distribution inferred with SeqDEFT (A) and the translational efficiencies inferred with VC regression (C). Every dot represents one of the possible 4^9^ possible sequences and is colored according to their inferred probability (A) or log(GFP) values (C). The inset represents the distribution of inferred sequence probabilities or log(GFP) values along with their corresponding color in the map. Inset in the upper right corner of (A) shows the relaxation times associated to the 20 most relevant Diffusion axes, showing that the first two Diffusion axes have much longer relaxation times than the rest. Sequences are laid out with coordinates given by these first two Diffusion axes and dots are plotted in the order of the 3rd Diffusion axis. (B) Two-dimensional histogram representing the relationship between the inferred sequence probabilities from their frequency in the *E. coli* genome and the estimated translational efficiencies inferred with VC regression from MAVE data. (D) Posterior distribution inferred by SeqDEFT for the scaled selection coefficient of specific mutations when introduced in two genetic contexts, UUAAGGAGC (grey) and UAAGGAGCA (black), representing a shift of the AGGAG motif by one nucleotide. Mutational effects are reported in units of scaled selection coefficients. (E) Posterior distribution inferred by SeqDEFT for the average scaled selection coefficient, relative to the average across all possible sequences, for genotypes containing the AGGAGG motif at positions separated by three nucleotides, along with their potential mutational intermediates. (F) Posterior distribution inferred with VC regression for the effects of specific mutations on log(GFP) when introduced in two genetic contexts, UUAAGGAGC (grey) and UAAGGAGCA (black), representing a shift of the AGGAG motif by one nucleotide.(G) Posterior distribution inferred with VC regression for the average log(GFP) of genotypes containing the AGGAGG motif at positions separated by three nucleotides, along with their potential intermediates. (E,G) Horizontal dashed lines represent posterior mean of the average phenotype across all possible sequences (grey) or wild-type sequences, given by the genomic sequences in (E) and AAGGAGGUG in (G) (black). Shaded areas represent the 95% credible intervals. (D-G) Points represent the maximum a posteriori (MAP) estimates and error bars represent the 95% credible intervals.

### Comparing sequence probability across different species

To investigate whether the structure of the genotype-phenotype map is the same across distant species, we performed the same analysis using 5’UTR sequences from 4,328 annotated genes in the genome of *B. subtilis*, whose most recent common ancestor with *E. coli* dates back to ∼2 billion years ago (Feng *et al*., 1997). We first found that the AG bias marking the location of the SD sequence in the 5’UTR is located approximately 2 bp further upstream from the start codon compared to its location in *E. coli* (Figure S6A), as previously reported (Hockenberry *et al*., 2018). We then extracted the 9 nucleotides sequences 6 bp upstream of the start codon and inferred the sequence probability distribution using SeqDEFT. The estimated log-probabilities were highly correlated with those obtained from the *E. coli* genome (Spearman *ρ* = 0.94, Figure S6B), but more importantly, the inferred genotype-phenotype map displayed a similar structure, with peaks corresponding to different binding registers of the aSD sequence and extended ridges connecting sets of sequences with overlapping binding sequences separated by 3 positions (Figure S6B). Overall, the probability distributions of the SD sequences are quantitatively very similar across distant species and show the same main qualitative features.

### Comparing sequence probability and functional measurements

We next compared the genotype-phenotype maps based on genomic sequences with the genotypephenotype map obtained with MAVE data. First, we directly compared the estimated sequence probability across the *E. coli* genome with the inferred translational efficiency from MAVE data (Figure 4B) for every possible sequence. We found a moderate non-linear relationship between these two independently inferred quantities (Spearman *ρ* = 0.55). Sequences with very low estimated probability (*P* < 10^−8^) consistently showed low translational efficiency (log(GFP) < 1.0), whereas sequences with high sequence probability (*P* > 10^−4^) had consistently higher but variable translational efficiencies (mean=1.84, standard deviation=0.63).

To investigate whether this modest degree of agreement is due to noise in the estimates for individual sequences or to having inferred qualitatively different genotype-phenotype maps, we applied the visualization technique to the empirical genotype-phenotype map inferred with VC regression (Figure 4C and S7). Despite the much more skewed phenotypic distribution of estimated translational efficiencies, this low-dimensional representation has essentially the same structure with isolated peaks corresponding to different distances of the SD motif to the start codon and extended ridges connecting sequences with SD motifs shifted by 3 positions separated along several Diffusion axes (Figure 4C and S8). In addition to the previous structure, we identify an additional extended ridge of functional sequences with sequences starting by GAG. This subsequence, together with the upstream G from the fixed genetic context in which the experiment was performed, forms a binding site for the aSD sequence. In contrast, the probability distribution of SD sequences was inferred from genomic sequences with different flanking nucleotides such that genotypes starting with GAG, on average, are not as functional. Thus, we can conclude that, despite showing only a moderate quantitative agreement, the two inference procedures using different types of data are able to recover genotype-phenotype maps with the same qualitative features and expected long-term evolutionary dynamics.

### Uncertainty quantification for genetic interactions and phenotypic predictions

Visualization of genotypephenotype maps based on our MAP estimates enabled the identification of key shared qualitative features and the genetic interactions underlying them. However, we can also evaluate the strength of evidence supporting these interactions by leveraging the uncertainty quantification capabilities of our Gaussian process models as implemented in *gpmaptools* e.g. we can compute the posterior distribution of the effects of specific mutations in different backgrounds. As an illustration of this strategy, we first validated the incompatibilities separating peaks in the SD genotype-phenotype map by computing the posterior distribution for mutational effects in the two backgrounds UUA**AGGAG**C (grey) and UA**AGGAG**CA (black), which contain the same AGGAG motif shifted by one position (Figure 4D). In absence of epistasis, mutational effects are expected to be exactly the same in the two genetic backgrounds. While this is true for some mutations e.g. C-5A (Figure 4D), the three mutations that allow shifting the SD motif one position up-stream in the UUA**AGGAG**C context (grey) i.e. A-10G, G-8A and A-7G, are strongly deleterious in that context, but beneficial when introduced in a UA**AGGAG**CA background (black, Figure 4D). Importantly, the posterior distributions are concentrated around the means, showing that the data strongly supports that mutations needed to shift the SD motif by one position are substantially deleterious in that speific context, creating the valleys that separate the main peaks of this genotype-phenotype map.

We next evaluated the evidence supporting the existence of the extended ridges connecting sequences with an SD motif shifted by 3 positions. To do so, we computed the posterior distribution for the average phenotype (scaled selection coefficient relative to the average fitness across all sequences) of genotypes containing two overlapping binding registers (NGGAGGAGN), only one (AGGAGGNNN and NNNAGGAGG), or none (NNNAGGNNN). Whereas NGGAGGAGN, AGGAGGNNN and NNNAGGAGG are highly functional, sequences with a central AGG have on average roughly half as large of a scaled selection coefficient than sequences containing full motifs in either or both registers (Figure 4E).

The posterior distributions for these same sets of genotypes and mutations estimated from sequence data from the *B. subtilis* genome (Figure S6D,E) and from VC regression analysis on MAVE data (Figure 4F,G) are largely concordant. The agreement between these independent data sources, together with uncertainty quantification in each case, provides strong support for a common landscape structure with distinct peaks at the different binding registers of the aSD sequence, connected by extended ridges of functional sequences linking registers offset by three nucleotides in aSD binding position.

### A biophysical model recapitulates the qualitative properties of empirical SD genotype-phenotype maps

Although the inferred genotype-phenotype map exhibited extensive epistasis, the visualization revealed that its complexity could be largely explained by a simple underlying mechanism in which the aSD sequence can bind at varying distances from the start codon. We hypothesize that this mechanism alone explains both the existence of isolated peaks and, together with the quasi-repetitive nature of the aSD sequence, the extended ridges. Moreover, despite our ability to estimate mutational effects in different contexts, inference of the actual binding preferences of the aSD from the data is hindered by the convolution of the effects of mutations on the binding affinities at different registers. To tackle these issues, we fit a simple mechanistic model, in which GFP protein abundance is linearly dependent on the fraction of mRNA bound by the aSD at thermodynamic equilibrium at different positions *p* relative to the start codon, where the binding energy Δ*G* of the aSD is an additive function of the sequence at that position *x*_*p*_ (Figure 5A, see Methods).

**Fig. 5.**
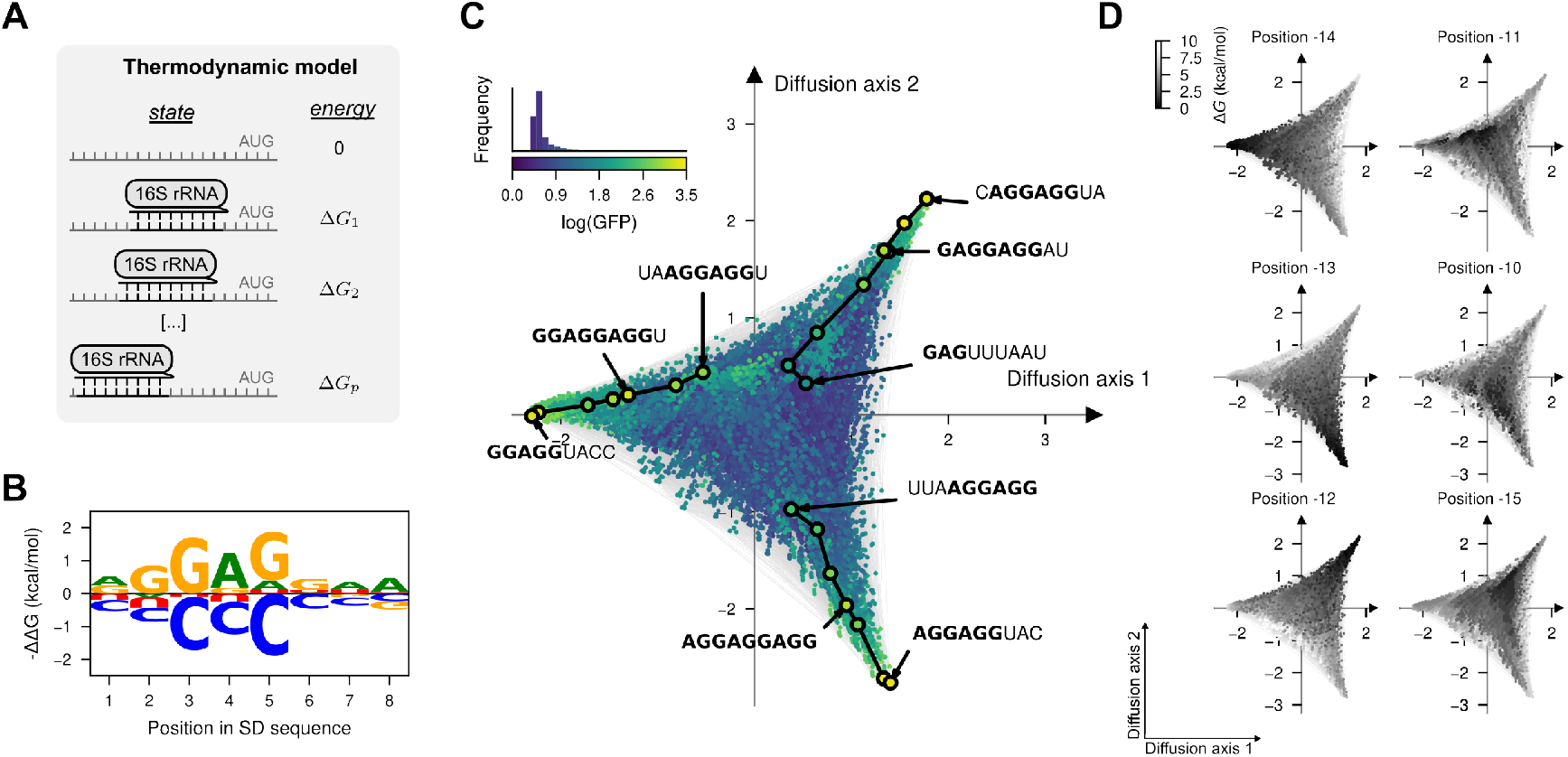
Biophysical model of sequence-dependent translational efficiency. (A) Thermodyanmic model of binding of the 16S rRNA 3’tail to the 5’UTR of mRNAs at different positions *p* relative to the start codon AUG. (B) Sequence logo representing the site-specific but register-independent allelic contributions to the binding energy, where the size of the letter represents the difference in binding energy to the average across nucleotides. (C) Visualization of the genotype-phenotype map that results from predicting the phenotype of every possible sequence under the inferred thermodynamic model. Every dot represents one of the possible 4^9^ possible sequences and is colored according to the predicted log(GFP). The inset represents the phenotypic distribution along with their corresponding color in the map. Sequences are laid out according to the first two Diffusion axes and dots are plotted in order according to Diffusion axis 3. (D) Visualization of the genotype-phenotype map under the inferred thermodynamic model representing the binding energies at positions −15 to −10 relative to the start codon showing that the peaks in the visualization correspond to the strongest binding at different positions and extended ridges correspond to sequences that are bound in two registers separated by 3 nucleotide positions. Binding energies are relative to the strongest binding sequence AGGAGGAA under the inferred model and are reported in units of kcal/mol assuming a temperature of 37ºC. Dots are plotted in reverse order of binding energy in the corresponding register.

We fit this biophysical model (Figure S9A) by maximum likelihood to the MAVE dataset and achieved good predictive performance in both training (*R*^2^ = 0.59, Figure S9B) and held-out sequences (*R*^2^ = 0.64, Figure S9C). Importantly, this model contains only 27 free parameters with clear biophysical interpretations e.g. in terms of mutational effects on binding energies. The model also includes a parameter *β*_0_ that specifies the background fluorescence in absence of aSD binding to the 5’UTR, which we estimated as 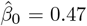.

This model allowed us to deconvolve the effects of mutations on binding at different registers and to infer the allele and position specific energetic contributions to binding (Figure 5B). As expected, the reverse complement of the aSD is the most stable binder, but different mutations have substantially variable effects in the binding energy. Not only do some positions have stronger energetic contributions in general (positions 2-5 within the SD sequence), but different missmatches with the aSD in the same position have different energetic effects e.g. A4G is only slightly destabilizing (ΔΔ*G* = 0.84 kcal/mol), whereas A4C is highly destabilizing ( the main structure of the genotype-phenotype map with isolated peaks and extended ridges corresponding to different registers of binding, as expected (Figure 5C). We can verify that the peaks correspond to different binding registers by computing the binding energy of every sequence at specific positions relative to the start codon and color the visualization by those energies (Figure 5D). Likewise, this representation also shows that the extended ridges of functional sequences correspond to sequences that are strongly bound at positions separated by three nucleotides. We next compared our thermodynamic model to an alternative thermodynamic model based on base-pair stacking interaction energies from classical RNA folding algorithms (Lorenz *et al*., 2011; Salis *et al*., 2009). Interestingly, our model achieved higher predictive accuracy (Figure S10A,B, *R*^2^ = 0.44) and more faithfully captured the overall structure of the empirical genotype-phenotype map (Figure S10C). These findings suggest that the SD:aSD interaction *in vivo* cannot be fully explained by RNA thermodynamics alone, highlighting the importance of molecular context in shaping RNA-RNA interactions. More broadly, this analysis shows how visualization can guide the construction of simplified, biophysically interpretable models that reproduce the key qualitative features of genotype-phenotype maps.

## Discussion

In this paper, we present *gpmap-tools*, an extensively documented software library with tools for the inference, visualization and interpretation of empirical genotype-phenotype maps containing arbitrarily complex higher-order genetic interactions. By providing a framework for the analysis of complex genetic interactions, *gpmap-tools* has the potential to reveal the simple qualitative properties of these complex mappings and to aid in development of biophysical and mechanistic hypotheses for these observed features.

The first step in this framework is the inference of the complete genotype-phenotype map comprising all possible sequences from either experimental MAVE data or sequence counts. Taking into consideration the noise in the data (due either to sampling noise or experimental error), *gpmap-tools* is capable of computing the high-dimensional posterior distribution over all possible genotype-phenotype maps under a variety of priors. This allows us to obtain the MAP estimate, that is, the most probable genotype-phenotype map given the observed data. However, in contrast to other expressive models able to capture complex genetic interactions, such as neural networks (Bryant *et al*., 2021; Gelman *et al*., 2021; Sethi and Zhou, 2024), our inference methods provide rigorous uncertainty quantification of phenotypes, mutational effects or any linear combination of phenotypic values. This is important, as it tells the user which phenotypic predictions, mutational effects or genetic interactions can be trusted and to what extent, given the data.

The second step in this framework is the interpretation of the inferred genotype-phenotype maps. *gpmap-tools* provides a powerful method for visualizing fitness landscapes (McCandlish, 2011) that allows exploratory data analysis, interpretation and comparison of complex datasets and models. Thus, rather than interpreting the results through an explicit parametric model allowing high-order genetic interactions (Poelwijk *et al*., 2016; Tareen *et al*., 2020; Faure and Lehner, 2024; Park *et al*., 2024; Faure *et al*., 2024) or descriptive statistics like the number of peaks or adaptive walks (Szendro *et al*., 2013; Ferretti *et al*., 2018; Papkou *et al*., 2023; Westmann *et al*., 2024b; Li and Zhang, 2025; Chattopadhyay *et al*., 2025), this method leverages the evolutionary dynamics on the genotype-phenotype map to highlight its main, potentially unexpected, qualitative features. Thus, we can use the visualization to generate hypotheses for how mutational effects change across genetic backgrounds and test these predictions by computing the corresponding background-dependent posterior distributions (Figure 4 and S6). Identifying the main features of the genotypephenotype map can be crucial for defining an appropriate mechanistic or biophysical model. Here, visualization of the Shine-Dalgarno landscapes allowed us to define a thermodynamic model in which the binding energy depends only additively on the sequence at each register, and to verify that this simple model recapitulated the main qualitative features of the landscape (Figure 5). Additionally, this technique enabled a detailed comparison of genotype-phenotype maps inferred with different methods and data sources. In contrast to broadly used metrics, like Pearson or Spearman coefficients, this method shows the extent to which different landscapes have the same structure and qualitative features. In this study, it showed that genotype-phenotype maps inferred from two distantly related species *E. coli* and *B. subtilis* (Figure 4 and S6), as well as from entirely independent data sources (MAVE experiments versus natural sequence data), exhibited strikingly similar structure despite only moderate quantitative agreement (Figure 4). Identifying consistent structures across different data types and sources is essential for linking experimentally measured landscapes to the evolutionary forces shaping regulatory sequences, given that true fitness values in natural populations are typically unknown. More broadly, our visualization technique enables comparison of genotype-phenotype maps across different classes of genetic elements, such as regulatory sequences, protein-protein interactions and enzymes, by revealing shared landscape features that may reflect similar evolutionary dynamics, despite differences in biological context.

The methods implemented in *gpmap-tools* scale to genotype-phenotype maps with millions of sequences by making several modeling assumptions, which also entail certain limitations. First, MEI and VC regression are phenomenological models. As such, they do not explicitly model global or non-specific epistasis that often arises from nonlinear dependencies between the underlying quantities affected by mutations and our measurements (Bloom, 2015; Otwinowski *et al*., 2018; Tareen *et al*., 2020; Tonner *et al*., 2022; Faure and Lehner, 2024). Instead, these models rely on learning the pervasive genetic interactions induced by these global nonlinearities to nonetheless make accurate phenotypic predictions. Second, SeqDEFT assumes that observed sequences are drawn independently from the underlying probability distribution. While this assumption may hold for a few specific regulatory sequences that are repeated many times along the genome of a single species e.g. the Shine-Dalgarno sequence or the 5’ splice site (Chen *et al*., 2021), it remains unclear how robust it is to the known challenge of using phylogenetically related sequences from widespread multiple sequence alignments of protein families (Hockenberry and Wilke, 2019; Rodriguez Horta and Weigt, 2021; Dietler *et al*., 2023). Third, both inference and visualization methods still require storing all possible sequences and their phenotypes in memory. The number of such sequences grows exponentially with sequence length, limiting the applicability of *gpmap-tools* to spaces of sequences of a constant and relatively short length (5 amino acids, 12 nucleotides, 24 biallelic sites). Despite these limitations, *gpmap-tools* provides a unique set of tools for studying the genotype-phenotype maps of short genetic elements. By combining nuanced analysis of epistasis, rigorous uncertainty quantification, and the capacity to infer landscapes containing millions of genotypes, it serves as a necessary stepping stone towards understanding the vastly larger genotypephenotype maps arising at the gene, protein, and genomewide scale.

## Methods

### Sequence diversity of the Shine-Dalgarno sequence

We downloaded the *E. coli* genome and annotation from Ensembl bacteria release 51, built on assembly version ASM160652v1, and *B. subtilis* assembly ASM904v1 from GeneBank. We extracted the 5’UTR sequence for every annotated gene using *pysam* (Li *et al*., 2009; Bonfield *et al*., 2021) and kept the 5,311 and 4,328 sequences, respectively, for which we could extract 20 bp upstream of the start codon without any ambiguous character ‘N’. These sequences were aligned with respect to the start codon and used for computing site-frequency logos using *logomaker* (Tareen and Kinney, 2020) and estimating the complex probability distribution using the *gpmap-tools* implementation of SeqDEFT (Chen *et al*., 2021). The MAP estimate was used to compute the coordinates of a low-dimensional representation assuming that the stationary distribution of the evolutionary random walk matches the estimated sequence probabilities by selecting a proportionality constant of *c* =1 and uniform mutation rates.

### Analysis of the experimental fitness landscape of the Shine-Dalgarno sequence

Phenotype data was computed from the processed data for independent replicates conducted in the dmsC genetic background as reported in the original manuscript (Kuo *et al*., 2020). The mean and standard error was computed for all the 257,565 measured sequences. We estimated a common measurement variance of 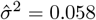 using genotypes measured across all 3 experimental replicates. The squared standard error for each genotype *i* was computed by dividing the overall experimental variance 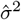 by the number of replicates *n*_*i*_ in which each sequence was measured 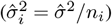.We kept 0.1% of the sequences as test set, and use the remaining sequences for fitting different models to infer the complete genotype-phenotype map while evaluating their performance on the held-out test data. We estimated the variance components from the empirical distance-correlation function and used them to define a Gaussian process prior for inference of the complete combinatorial landscape containing all 4^9^ genotypes, taking into account the known experimental variance 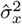 for every sequence. We also computed the posterior mean and variances across all test sequences to assess the accuracy of the predictions and the calibration of the posterior probabilities in held-out data. We used the MAP estimate to compute the coordinates of the visualization assuming several different average values of log(GFP) under the stationary distribution that ranged from 1 to 2.5 (Figure S7). An average log(GFP) of 2 at stationarity was selected and used for all subsequent visualizations, similar to our MAP estimate of a log(GFP) of 2.03 for the wild-type reference.

### Thermodynamic model of the Shine-Dalgarno genotype-phenotype map

We assume that translation is limited by the initiation step, which is itself modulated by the binding of the 16S rRNA to the 5’UTR of the mRNA, where we assume that the mRNA concentration is independent of the identity of the Shine-Dalgarno sequence. Binding and dissociation are assumed to be much faster than the rate at which translation is effectively initiated, so that the protein abundance is proportional to the fraction of mRNA bound by the 16S rRNA across all registers *p* at thermodynamic equilibrium, where we assume that binding occurs in at most one register at a time. The fraction of mRNA bound at thermodynamic equilibrium depends on the binding energy Δ*G* of the 16S rRNA to the mRNA to the sequence *x*_*p*_ starting at each position *p*, the temperature, which is assumed to the 37ºC (310K), and the universal gas constant *R* = 1.9872 × 10^−3^ kcal/mol K^−1^. The overall GFP concentration for a sequence *x* depends on the fraction of bound mRNA and the translation rate when bound *β*:

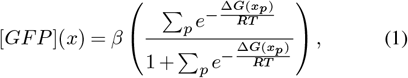

where Δ*G*(*x*_*p*_) is the energy of binding of the 16S rRNA to the 8-nucleotide subsequence *x*_*p*_ at position *p*. The binding energy Δ*G* is independent of the position *p* at which binding occurs relative to the start codon and depends additively on the sequence *x*_*p*_ alone given by Δ*G*(*x*_*p*_)= Δ*G*_0_ + ∑_*i*_ ∑_*c*_ *x*_*p*_(*i, c*)ΔΔ*G*_*ic*_, where *x*_*p*_(*i, c*) takes value 1 if se-quence *x*_*p*_ has allele *c* at position *i* and 0 otherwise, Δ*G*_0_ represents the average binding energy across every possible sequence and ΔΔ*G*_*i,c*_ is the energetic contribution of allele *c* at position *i*, subject to the constraint _*c*_ ∑_*c*_ΔΔ*G*_*i,c*_ =0 across all positions *i*. In order to incorporate the effect of mutations in binding registers spanning both fixed and variable regions of the sequence, we extended the variable 9 nucleotide sequences with the fixed upstream and downstream sequences CCG and UGAG from the dmsC genetic context.

Following previous work (Kuo *et al*., 2020), we assume that occupancy at thermodynamic equilibrium is low so tha 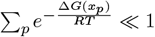, and thus [*GFP*] 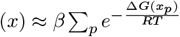. We also model a background fluorescence signal *β*_0_ due to cells auto-fluorescence in the GFP channel even in absence of GFP, which is independent of the variable 5’UTR sequence in the experiment. Finally, we consider that experimental errors lie on the log-scale, such that the measured log(GFP) *y* for sequence *x* is observed with known noise variance 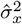 and an extra or uncharacterized variance *σ*^2^ under a Gaussian likelihood function given by

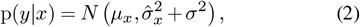

where *μ*_*x*_ is the expected log(GFP) under the model given by

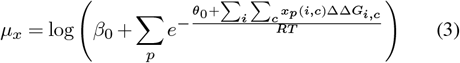

and *θ*_0_ = *RTβ* + Δ*G*_0_. We used PyTorch to encode the model and used the Adam optimizer with a learning rate of 0.02 for 1500 iterations, while monitoring for convergence (Figure S9A), to find the maximum likelihood estimates of the model parameters.

Additionally, we fit a 4-parameter calibration model using ensemble binding energies Δ*G*_*x*_ computed with a thermodynamic model of RNA folding and interaction (Lorenz *et al*., 2011), where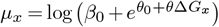. Specifically, we used RNAcofold v2.4.9 with -p0 option for computing the binding energy of each SD variant, embedded between the CCG and UGAG flanking sequences, with the anti-SD sequence ACCUCCU across all possible binding configurations. Several candidate anti-SD sequences, ranging from 5 to 9 nucleotides, were tested; ACCUCCU was selected due to the higher predictive power of GFP abundance of the resulting model.

## Code availability

*gpmap-tools* is an open-source library with source code available at https://github.com/cmarti/gpmap-tools. It is thoroughly documented with several tutorials and explanations of the provided functionalities at https://gpmaptools.readthedocs.io. Code to reproduce the analyses of the Shine-Dalgarno landscapes is available at https://github.com/cmarti/shine_dalgarno.

## Acknowledgements

We thank Bryan Gitschlag, Álvaro Serrano-Navarro, Víctor Jiménez-Jiménez and Alejandra Laguillo-Diego for providing feedback during the preparation of this manuscript. CMG and DMM were supported by NIH grant R35GM133613, JBK and DMM were supported by NIH grant R01HG011787, JBK was supported by NIH grant R35GM133777, and CMG, JBK, and DMM were supported by additional funding from the Simons Center for Quantitative Biology at Cold Spring Harbor Laboratory. JZ was supported by NIH grant R35GM154908. WCC was supported by the National Science and Technology Council of Taiwan, R.O.C., under Grant No. NSTC 111-2112-M-194-008-MY3.

This work was performed with assistance from the US National Institutes of Health Grant S10OD028632.

## Supplementary Figures

**Fig. S1.**
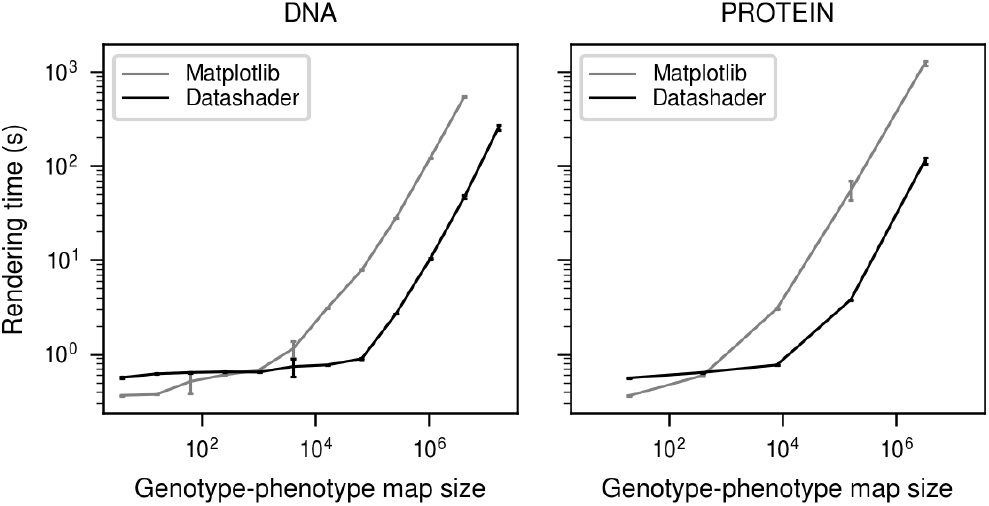
Visualization rendering times using two different back-end libraries for plotting as a function of the size of DNA and protein genotype-phenotype maps.

**Fig. S2.**
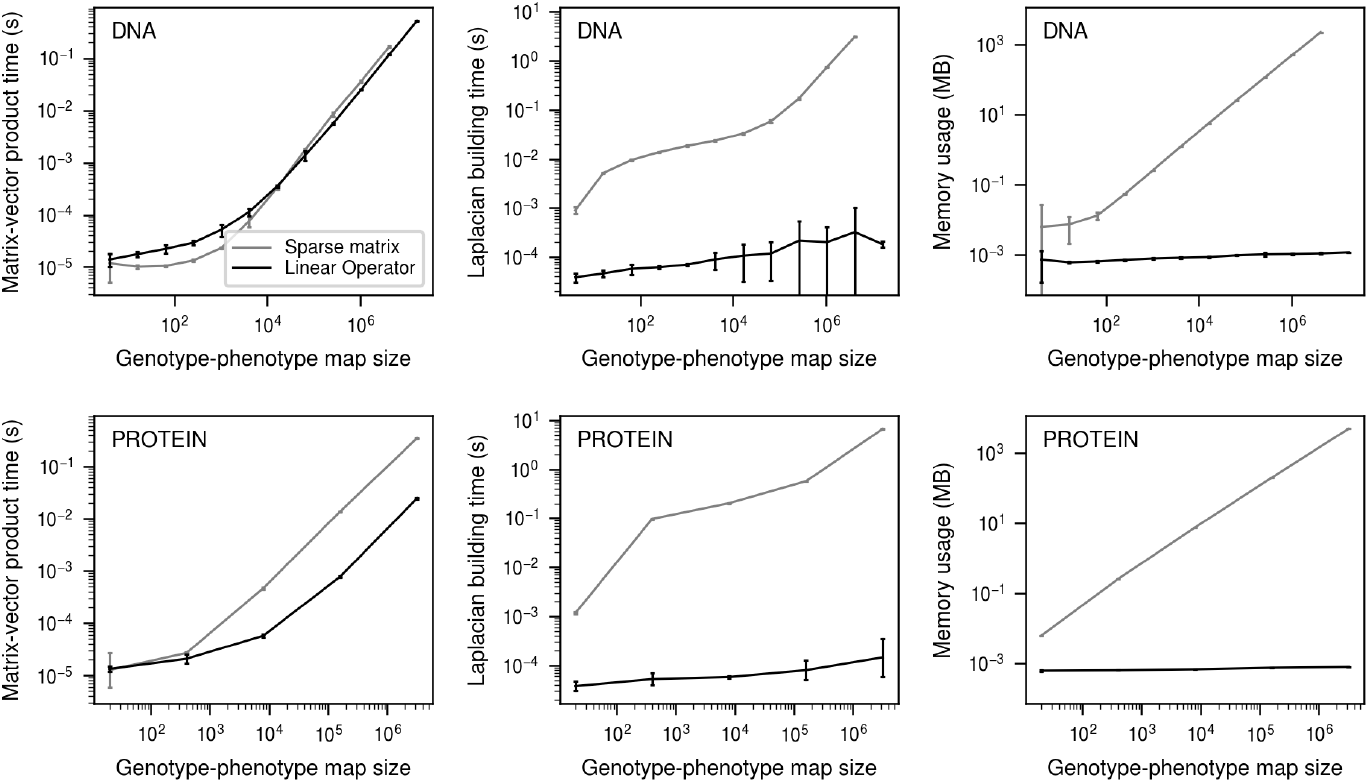
Comparison of the running times and memory requirements for computation of matrix-vector products with the Laplacian of the Hamming graph using our new Linear Operator or our previous sparse matrix formulation.

**Fig. S3.**
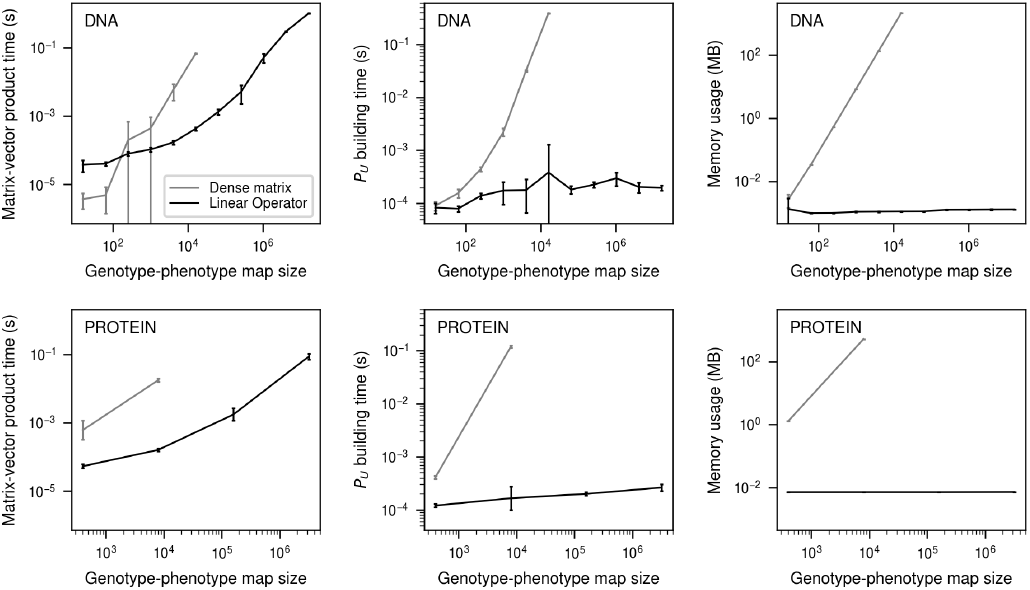
Comparison of the running times and memory requirements for computation of *P*_*U*_ matrix-vector products using our Linear Operator or the corresponding dense matrix.

**Fig. S4.**
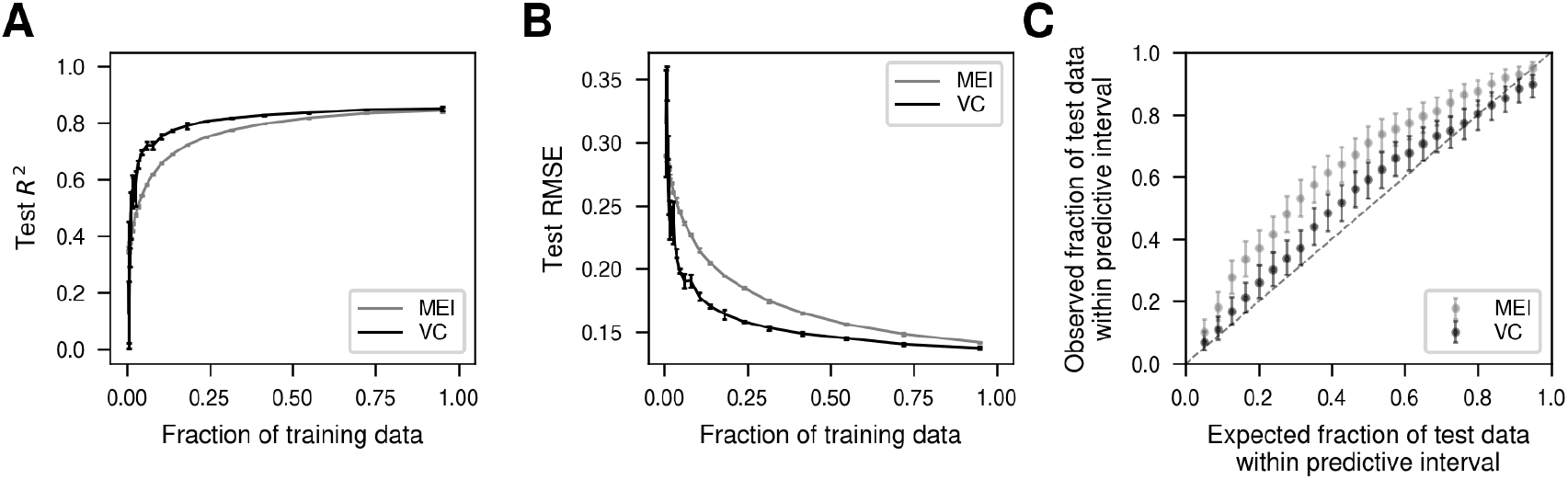
Predictive performance of Minimum Epistasis Interpolation (MEI) and Variance Component (VC) regression in held-out data. (A,B) Model predictive performance measured by the *R*^2^ (A) and RMSE (B) in held-out data as a function of the fraction of data used for training and phenotypic prediction. Error bars represent the standard deviation over 3 independent subsets of sequences used for training at each proportion. (C) Evaluation of the models calibration by comparing the expected fraction of times a predictive interval will contain the real phenotypic value compared to the fraction of times it actually contained the measured phenotype across 274 test data points. Error bars represent the 95% Jeffreys confidence interval for the estimated fraction of data points laying within the corresponding predictive interval. Diagonal dashed gray line shows the expectation under perfect model calibration.

**Fig. S5.**
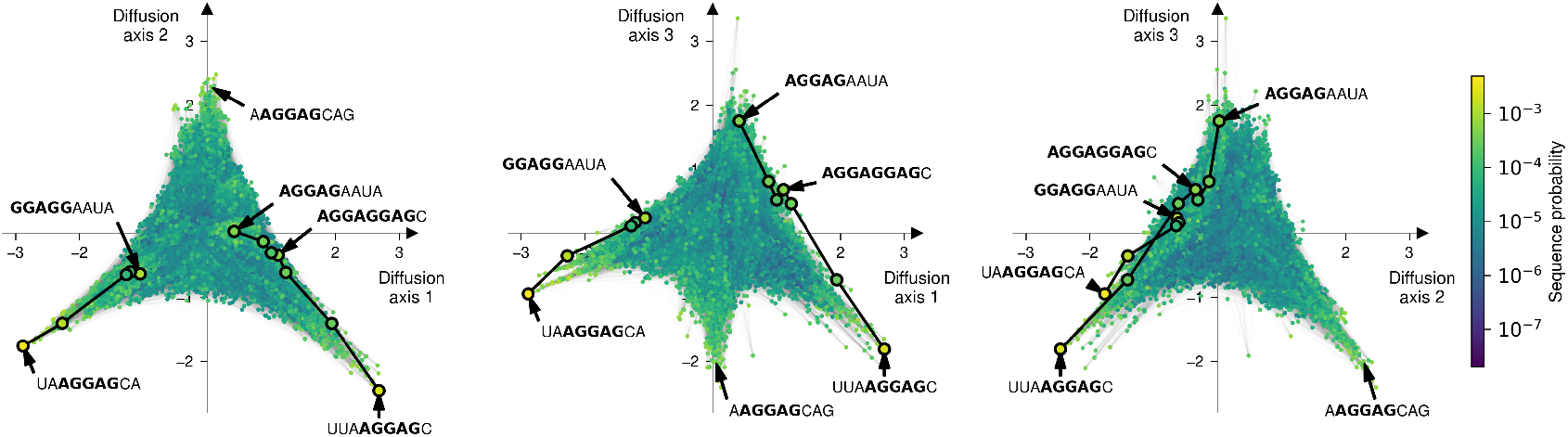
Low-dimensional representation of the Shine-Dalgarno probability distribution inferred with SeqDEFT along Diffusion axes 1, 2 and 3. Every dot represents one of the possible 4^9^ possible sequences and is colored according to their inferred probability. Sequences are laid out according to the indicated Diffusion axes and dots are plotted in order according to the missing Diffusion axis.

**Fig. S6.**
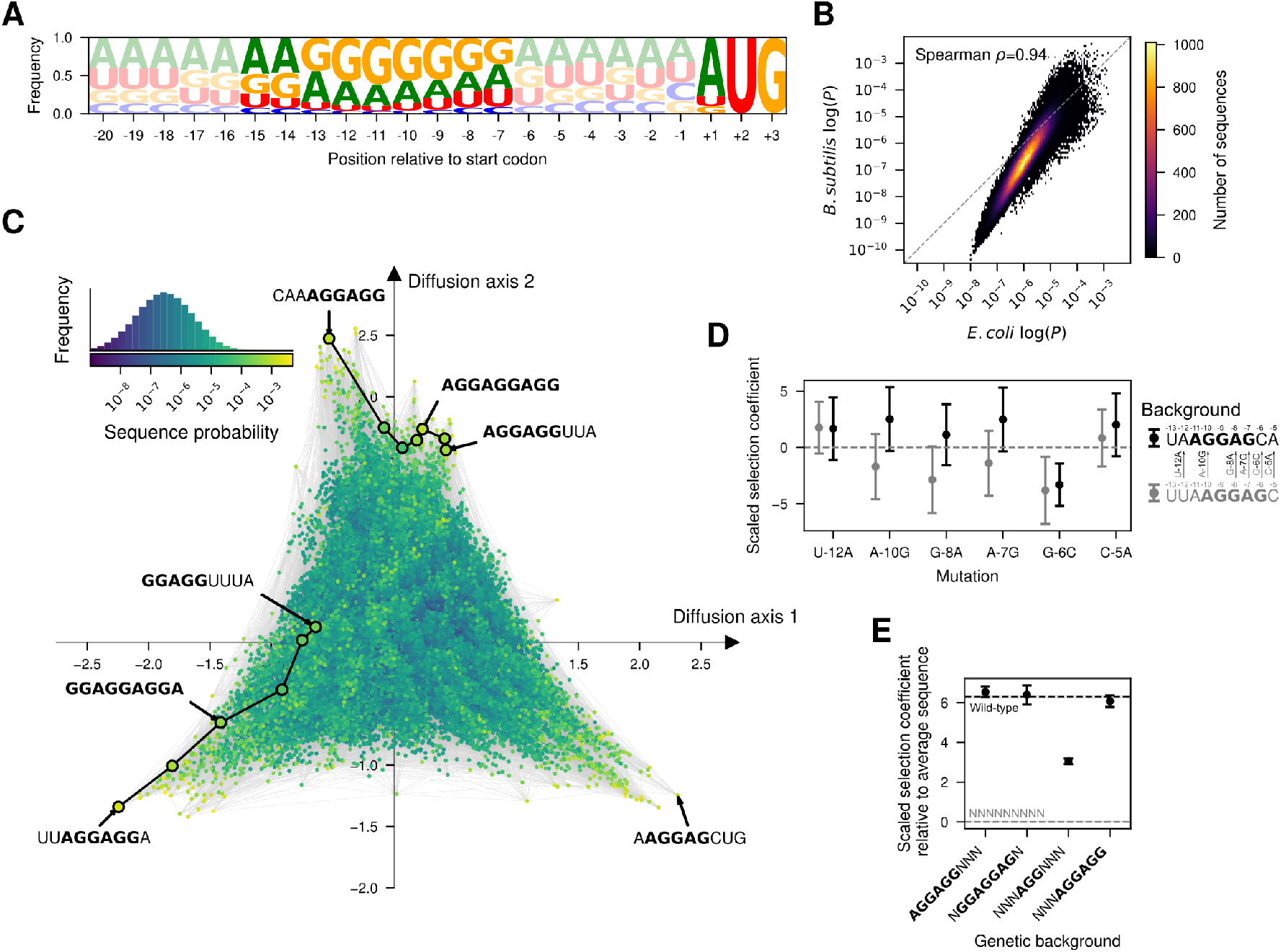
The structure of the genotype-phenotype map inferred from *B. subtilis* is conserved. (A) Sequence logo representing the site-specific allele frequencies of 4,328 5’UTRs in the *B. subtilis* genome aligned with respect to the annotated start codon. The start codon and the 9 nucleotide sequences 6 bases upstream are highlighted to emphasize the most relevant cis-regulatory sequences for translation initiation. (B) Two-dimensional histogram representing the relationship between the inferred sequence probabilities from their frequency in the *E. coli* and *B. subtilis* genomes. (C) Low-dimensional representation of the Shine-Dalgarno probability distribution inferred with SeqDEFT. Every dot represents one of the possible 4^9^ possible sequences and is colored according to its inferred probability. The inset represents the distribution of inferred sequence probabilities along with their corresponding color in the visualization. Sequences are laid out according to the first two Diffusion axes and dots are plotted in order according to the 3rd Diffusion axis. (D) Posterior distribution inferred by SeqDEFT for the scaled selection coefficient of specific mutations when introduced in two genetic contexts, UUAAGGAGC and UAAGGAGCA, representing a shift of the AGGAG motif by one nucleotide. Note that estimated mutational effects, except for G-6C, are largely compatible with those estimated from the *E. coli* genome shown in Figure 4D in the UAAGGAGCA context. (E) Posterior distribution inferred by SeqDEFT for the average scaled selection coefficient, relative to the average across all possible sequences, for genotypes containing the AGGAGG motif at positions separated by three nucleotides, along with their potential mutational intermediates. Horizontal dashed lines represent posterior mean of the average phenotype across all possible sequences (grey) or wild-type genomic sequences (black). Shaded areas represent the 95% credible intervals. (D,E) Points represent the maximum a posteriori (MAP) estimates and error bars represent the 95% credible intervals.

**Fig. S7.**
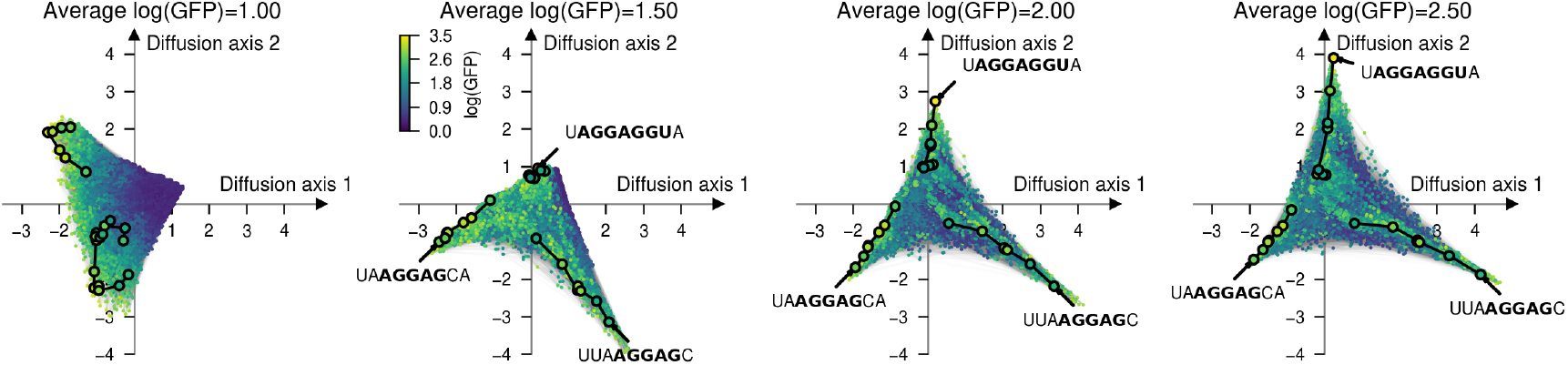
Low-dimensional representation of the Shine-Dalgarno genotype-phenotype map inferred with VC regression from MAVE data along Diffusion axes 1 and 2 as a function of the assumed average log(GFP) at the stationary distribution (as determined by tuning the strength of selection parameter *c*). Every dot represents one of the 4^9^ possible sequences and is colored according to its inferred log(GFP) values. Dots are plotted in order according to Diffusion axes 3.

**Fig. S8.**
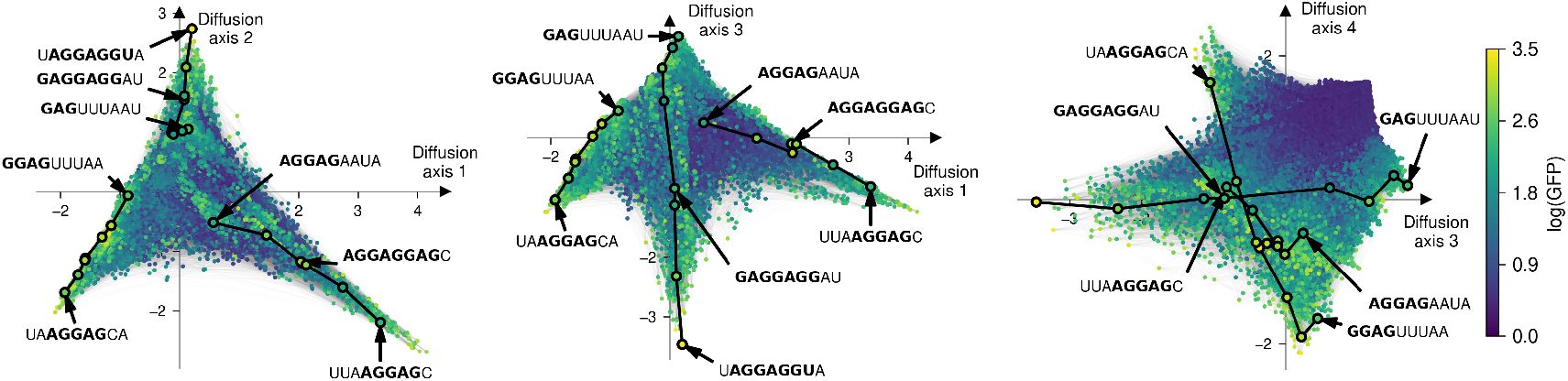
Low-dimensional representation of the Shine-Dalgarno genotype-phenotype map inferred with VC regression from MAVE data along Diffusion axes 1, 2, 3 and 4. Every dot represents one of the possible 4^9^ possible sequences and is colored according to its inferred log(GFP) value. Sequences are laid out according to the indicated Diffusion axes and dots are plotted in order according to Diffusion axes 3, 2 and 1, respectively for each panel.

**Fig. S9.**
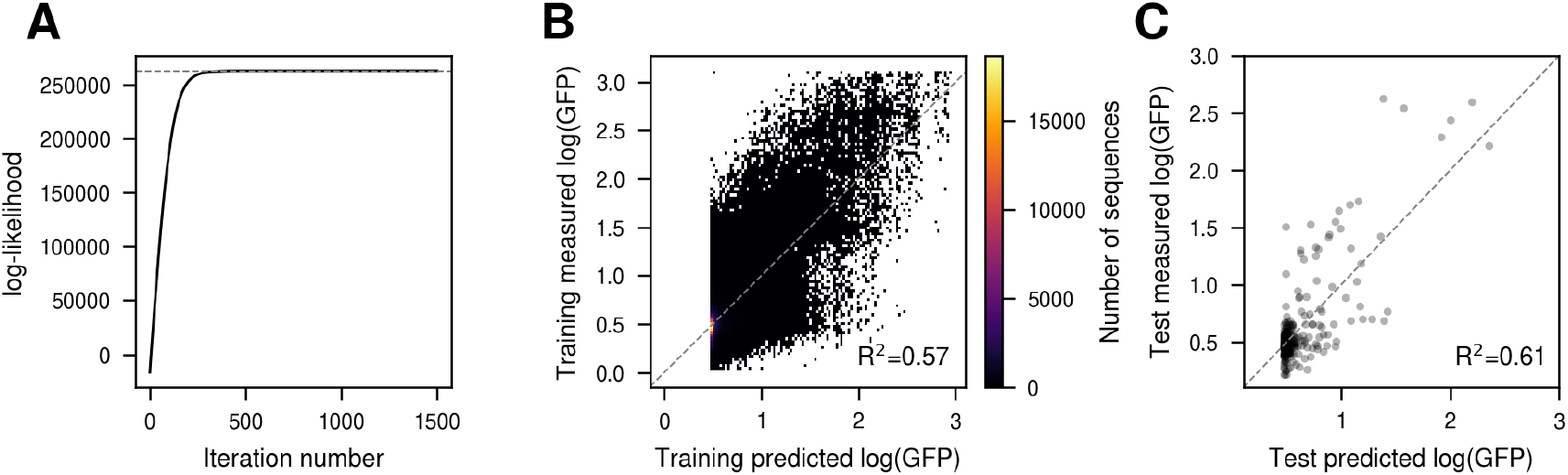
Fitting a thermodynamic model to the Shine-Dalgarno genotype-phenotype map using MAVE data. (A) Training curve showing the evolution of the log-likelihood as a function of the number of iterations of the Adam optimizer. (B) Comparison of measured log(GFP) in the training data with the predicted values under the estimated thermodynamic model. (C) Comparison of measured log(GFP) in the test data with the predicted values under the estimated thermodynamic model.

**Fig. S10.**
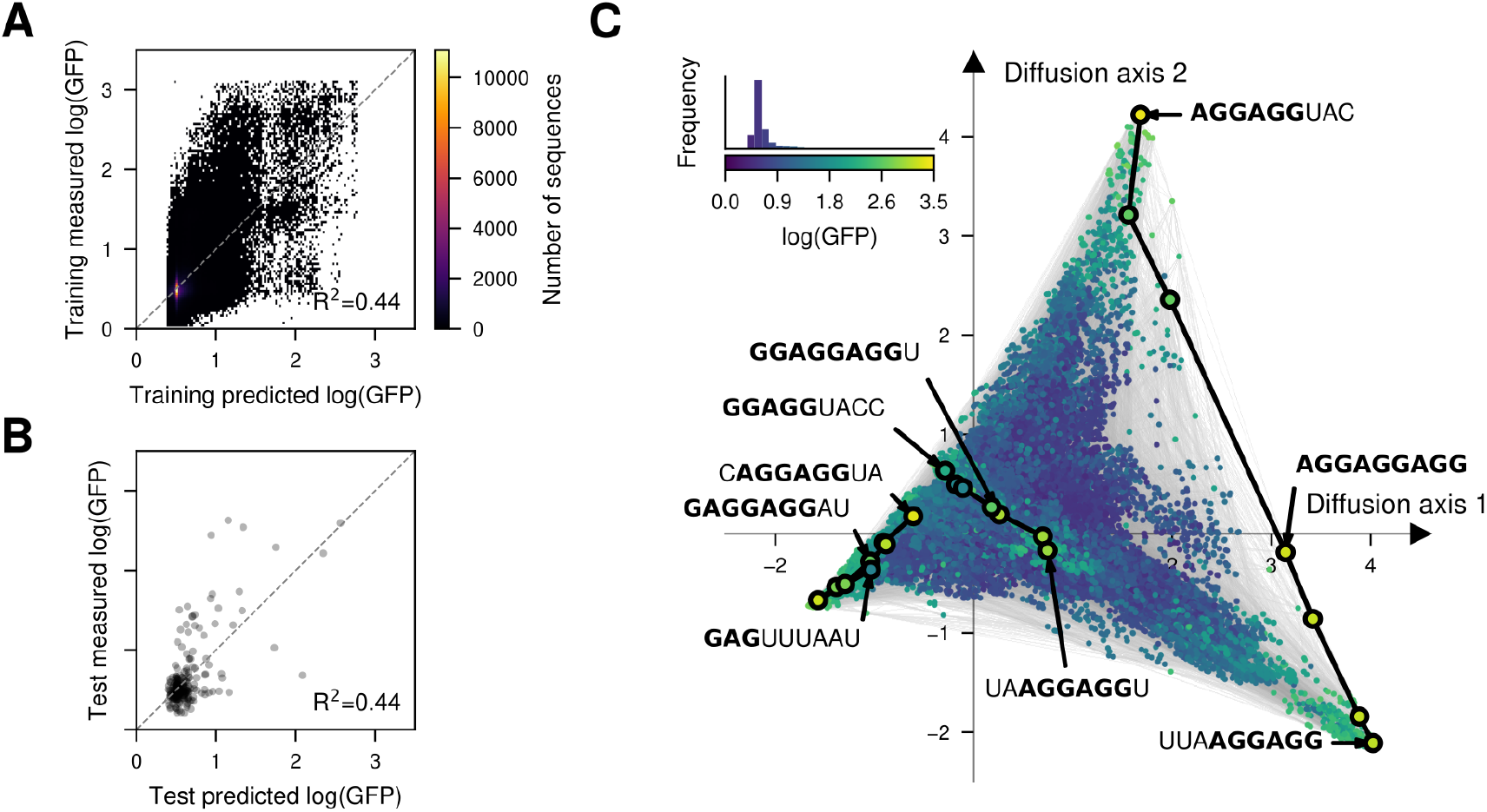
A general thermodynamic model for RNA folding for predicting MAVE data for the Shine-Dalgarno sequence. (A,B) Comparison of measured log(GFP) in the training (A) and test data (B) with the predicted values under the calibration model using RNAcofold ensemble binding energies with the anti-SD sequence. (C) Visualization of the genotype-phenotype map that results from predicting the phenotype of every possible sequence RNAcofold ensemble binding energies to the anti-SD sequence. Every dot represents one of the possible 4^9^ possible sequences and is colored according to the predicted log(GFP). The inset represents the phenotypic distribution along with their corresponding color in the map. Sequences are laid out according to the first two Diffusion axes and dots are plotted in order according to Diffusion axis 3.

## Supplementary Information

### A. Properties of the *P*_*U*_ projection operators

Let *f* represent a genotype-phenotype map on the space of sequences with a single site and *α* different alleles. The function *f* can be projected into a subspace spanned by any vector *b* using the orthogonal projection matrix 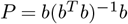. Letting *V*_*con*_ be the constant subspace spanned by the *α*-dimensional vector of ones 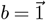, the orthogonal projection matrix into *V*_*con*_ is thus given by 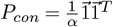. Let *V*_*add*_ be the orthogonal subspace (*V*_*con*_ ⊥ *V*_*add*_), defined by its projection matrix 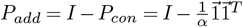. Thus, for any pair of sequences 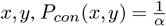 and

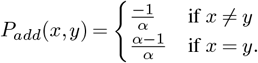

For a genotype-phenotype map *f* in the space of sequences of length *ℓ*, these elementary subspaces can be combined through tensor products into 2^*ℓ*^ different 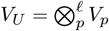 subspaces defined by the set of sites *U*, such that *V*_*p*_ = *V*_*add*_ for *p* ∈ *U* and *V*_*p*_ = *V*_*con*_ for *p* ∉*U*. Thus, the projection operator into the subspaces defined by *U* are obtained through the Kronecker product 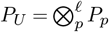, the elements of which are

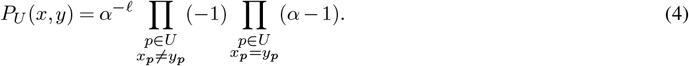

It is easy to show that the resulting subspaces *V*_*U*_ are orthogonal to each other using the mixed product property of the Kronecker product (*A* ⊗ *B*)(*C* ⊗ *D*) = (*AC*) ⊗ (*BD*). As a consequence, if either *AC* = 0 or *BD* = 0, then *A* ⊗ *B* and *C* ⊗ *D* are orthogonal. If we consider two subspaces defined by different subsets of sites *U* and *U* ^t^, there will be at least one position *p* at which the two site-specific elementary subspaces are orthogonal to each other, i.e. 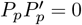. Consequently, 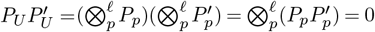.

Next, we consider the subspaces defined by the direct sum of subspaces *V*_*U*_ defined by exactly *k* sites 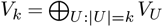 and derive the corresponding projection operator *P*_*k*_:

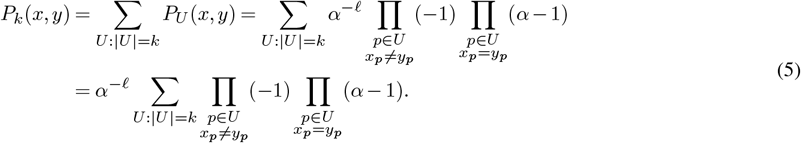

We note that the elements in the sum are obtained by multiplying the factors (*α*−1) and (−1) together a total of *k* times. These products can take only *k* +1 possible values: (−1)^*q*^ (*α*−1) ^*ℓ*−*q*^ for *q* = 0, 1,…,*ℓ*, where *q* represents the number of missmatches between sequences *x* and *y* within sets of sites *U*. Therefore, *P*_*k*_ can be expressed by:

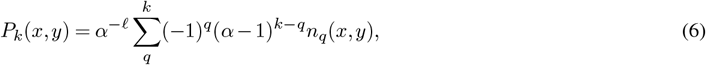

where *n*_*q*_(*x, y*) is the number of times each unique value appears when summing through the corresponding *P*_*U*_ matrices. Because we are summing over all possible *U* of the same size, *n*_*q*_(*x, y*) does not depend on the specific sites or alleles at which *x* and *y* differ, but only on the Hamming distance between them *d*(*x, y*). Specifically *n*_*q*_(*x, y*) is obtained by multiplying the number of ways to select *q* sites within the set of *d*(*x, y*) different sites by the number of ways to select the remaining *k* − *q* sites within the set of *f* − *d*(*x, y*) matching sites:

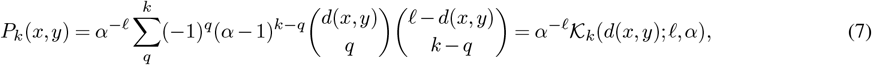

where 𝒦_*k*_(*d*(*x, y*); *ℓ, α*) denote the Krawtchouk polynomials (Stadler *et al*., 1994; Zhou *et al*., 2022). Note that this expression corresponds to the projection operator *P*_*k*_ into the subspace of pure *k*-th order interactions. Therefore, this shows that this *k*-th order interaction subspace *V*_*k*_ can be decomposed into smaller orthogonal subspaces *V*_*U*_ corresponding to pure *k*-th order interactions involving specific subsets of sites.

It is also easy to show that the columns of *P*_*U*_ are in the *k*-th eigenspace of the graph Laplacian by taking its projection and using the orthogonality properties of the *P*_*U*_ :

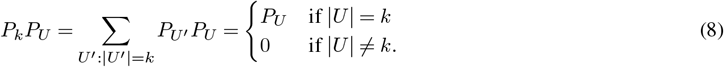

Using the same argument, we can show that 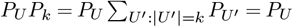 when |*U* | = *k* and 0 otherwise.

### B. Variance explained by interactions among sets of sites

In section A, we introduced the projection matrix *P*_*U*_. This enables decomposing a genotype-phenotype map *f* into its 2^*ℓ*^ components *f*_*U*_, each representing pure *U* -order interactions among sites in a set |*U*. Here, we present two coarse-grained views of these components by quantifying the percentage variance explained by genetic interactions involving specific sites in a genotype-phenotype map *f*, where the total variance is given by

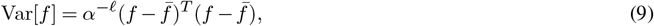

Where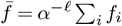. In the first approach, we consider the variance explained by interactions of order *k* involving each site *p* in the sequence:

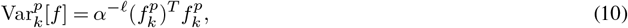

where

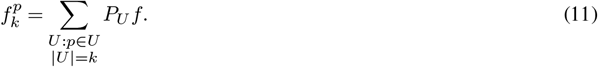

The values 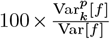 for every combination of *k* and *p* can be arranged in an *ℓ* × *ℓ* -dimensional matrix to reveal which sites contribute most to variance, and how much through additive, low-order or higher-order genetic interactions.

In the second approach, we consider the variance explained by interactions of order *k* =2 and *k*> 2 involving pairs of sites *p, q*:

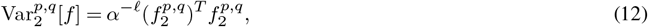

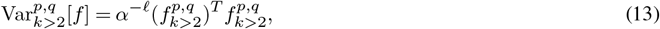

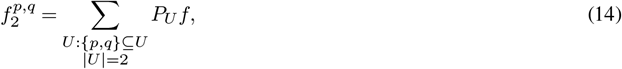

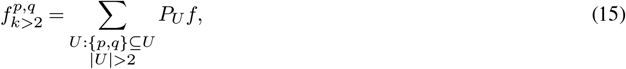

These values can be summarized in an *ℓ* × *ℓ* -dimensional matrix *M*

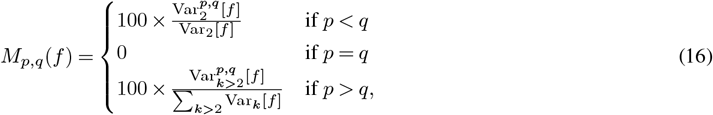

Where 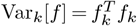. This matrix reveals the interaction patterns between pairs of sites, distinguishing pairwise from higher-order effects.

### C. Linear operator for Kronecker products

In section A, we describe the projection matrix *P*_*U*_ that projects a function *f* into the subspace corresponding to pure interactions between sites in a set *U*. Here we describe an efficient method for calculating the product *P*_*U*_ *f* by noting that *P*_*U*_ can be written as a Kronecker product of *ℓ* matrices 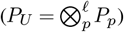.

Let *A* be an arbitrary matrix obtained through an *f*-Kronecker product, i.e. 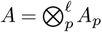 with entries given by *A*(*x, y*) = Π_*p*_ *A*_*p*_ (*x*_*p*_,*y*_*p*_) and **B** ∈ ℝ^(*α*×*α*×…×*α*)^ be a tensor with *ℓ* dimensions such that 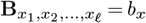 for any vector 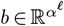. Thus, any matrix vector product *Ab* can be computed without explicitly constructing *A* using tensor dot products as follows:

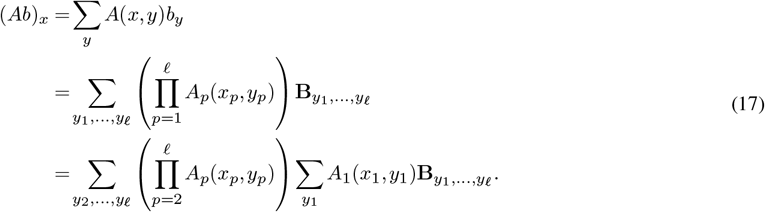

Let 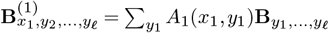 and repeat the same operation:

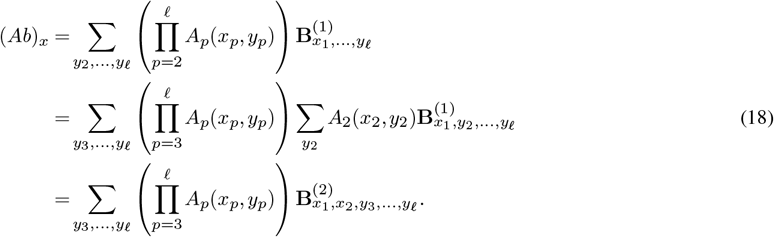

Thus, it is easy to see that (*Ab*)_*x*_ corresponds to 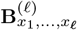 under the general recursion

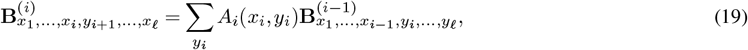

Where 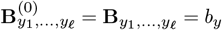. Note that each of the *ℓ* steps in the recursion can be computed efficiently as a tensor dot product between an *α* × *α* matrix and an *α*_1_×_2_*α*×···×*α*_*ℓ*_ tensor requiring *α*^*ℓ*−1^ *α*^2^ operations. The total number of required operations, *ℓα*^*ℓ*+1^, is much smaller than the *α*^2*ℓ*^ operations required for naively computing *Ab*, but more importantly, this strategy reduces memory requirements from the prohibitive scaling with *α*^2*ℓ*^ for storing *A* to only *α*^*ℓ*^ for storing **B**, enabling practical computation for *α*^*ℓ*^ in the order of millions.

### D. Linear operator for Laplacian of the Hamming graph

The space of possible sequences of length *ℓ* and *α* different alleles can be represented by a Hamming graph, in which nodes represent genotypes and edges represent single point mutations. The Laplacian matrix *L* of this graph is given by

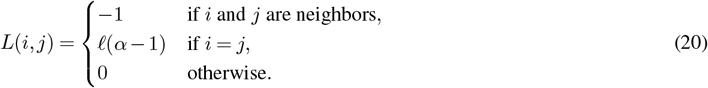

This matrix is sparse and can be stored in Compressed Sparse Row (CSR) format to efficiently compute matrix vector products. Despite the sparsity, there are still *α*^*ℓ*^ (1 + *ℓ*(*α ™* 1)) non-zero entries. For a the space of sequences of length 9 with 4 alleles with 64 bits floating point values, only storing the non-zero entries would require about 450MB. Here, we develop a matrix-free function to compute matrix-vector products with the Laplacian matrix *Lb* by leveraging the highly regular structure of this matrix and tensor broadcasting, resulting in memory requirements that scale only with the size of sequence space *α*^*ℓ*^. In particular, we can express *Lb* as

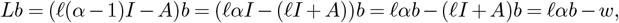

where *A* is the adjacency matrix and *w* is the product (*ℓI* + *A*)*b*. For a fixed choice of *b*, let **B** ∈ ℝ^(*α*×*α*×…×*α*)^ be a tensor with *ℓ* dimensions such that 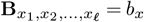, where *x* represents a sequence and *x*_*i*_ the allele at position *i*. Then for the same choice of *b*, the tensor **W** having elements 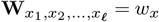 can be easily computed in tensor form using broadcasting by using the trick

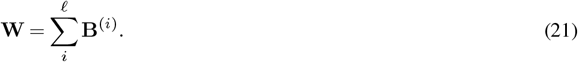

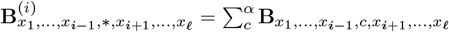, where the ‘*’ character indicates broadcasting, i.e. all characters at position *i* lead to the same value. Thus, this can be efficiently computed by summing the entries of tensor **B** over axis *i*. We can then use **B** and **W** to calculate *w* = (*ℓI* + *A*)*b* as:

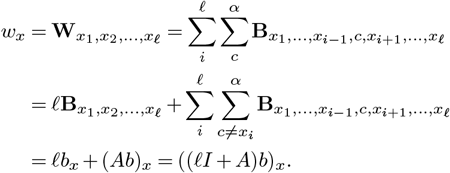

### E. Minimum epistasis interpolation solution and posterior distribution

In this section, we derive the minimum epistasis interpolation solution as the maximum a posteriori (MAP) estimate of a Gaussian process model under a prior distribution on local epistatic coefficients. Let us consider a complete genotype-phenotype map given by an *α*^*ℓ*^-dimensional vector *f* and define an improper prior distribution defined by the precision matrix 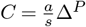, such that log p(*f*) ∝− *f* ^*T*^ *C f*.

Assuming we know exactly the phenotypes *f*_*x*_ for a subset of sequences *x*, we aim to compute the posterior probability of the phenotypes *f*_*z*_ at all remaining unobserved sequences *z* given by p(*f*_*z*|_*f*_*x*_). Let us define the joint log-probability distribution over [*f*_*x*_, *f*_*z*_] given by

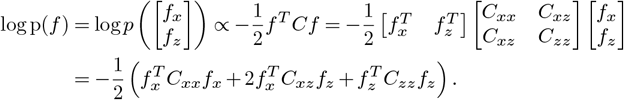

Since *C*_*zz*_ is a principal submatrix of the positive semidefinite matrix *C*, it is also positive semidefinite for any *z*, and thus log p(*f*_*z*_|*f*_*x*_) is convex. We can now take the gradient 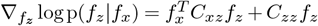 and find the MAP 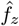 as the solution to the equation 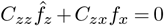, which is unique if and only if the corresponding *P* − 1-th order model is uniquely determined (Zhou and McCandlish, 2020) and is given by 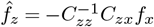. In this case, it is easy to verify that the posterior covariance is given by 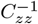 as

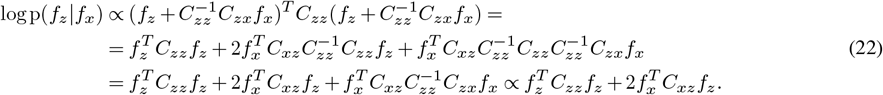

In the case in which 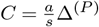, the posterior mean 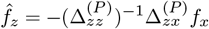 is independent of *a*. The posterior covariance, on the other hand, is independent of *f*_*x*_, but depends on *a* and the pattern of observations. Thus, if we want to compute the variance of the phenotypic predictions of specific sequences, we define the *a*^∗^ such that the expected sum of squared local epistatic coefficients 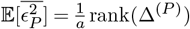 under the prior (Chen *et al*., 2021) matches the one under the posterior mean given by 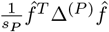.

### F. Posterior distribution computation with the cost matrix

Given a set of *n* measurements *y* in a subset of sequences *x* with measurement variances arranged along the diagonal of an *n* × *n* matrix *D*, we aim to obtain the complete genotypephenotype map represented by the *α*^*ℓ*^-dimensional vector 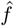 that maximizes the posterior probability of *f* given the observations *y*, i.e. we wish to find

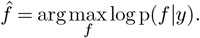

We begin by defining an *α*^*ℓ*^ × *n* matrix *X* relating the points in the complete space with the *n* observed values, such that *f*_*x*_ = *X*^*T*^ *f*

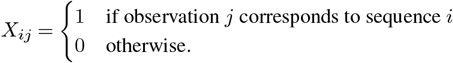

Using *X*, we can then write the posterior log-probability as a function of *f* :

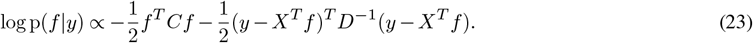

We can then expand this expression to separate factors that depend on *f* from those that depend only on the data *y*.

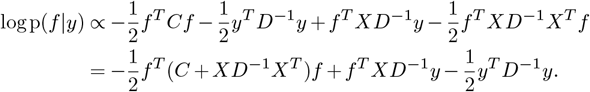

We next take the gradient with respect to *f*

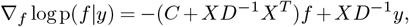

and solve for 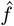 by setting ∇_*f*_ log p(*f* |*y*)= 0, which yields

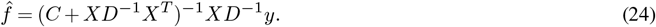

If *C* is invertible, then we can define a kernel matrix *K* = *C*^−1^ over the complete genotype-phenotype map and verify that this is equivalent to the classical solution for the posterior mean of a Gaussian process model using Woodbury’s identity. Specifically, for a subset of sequences *z* and the *α*^*ℓ*^ by |*z*| matrix *Z* defined by:

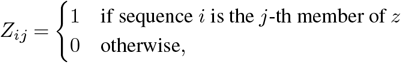

we find that

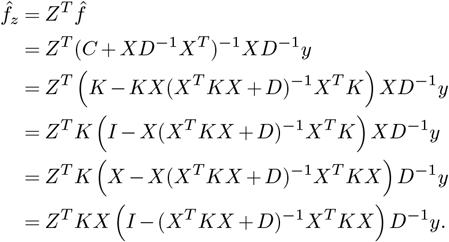

Then we can use the fact that *I* = (*X*^*T*^ *KX* + *D*)^−1^(*X*^*T*^ *KX* + *D*) to obtain the identity:

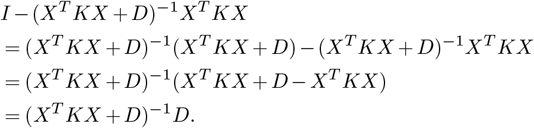

Substituting this identity into our previous expression for 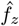, we now recover the classical maximum a posteriori solution for Gaussian process regression

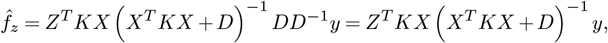

as desired.

Turning to the covariance matrix for the posterior, knowing that the posterior distribution is multivariate Gaussian implies that the posterior covariance matrix is given by the inverse of the Hessian matrix of the log-posterior probability. Thus the covariance matrix of the posterior distribution is given by:

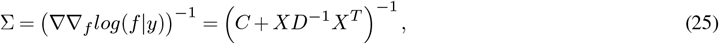

which we note depends on our observations only through the pattern of observed sequence as encoded in *X* and not on the observed phenotypes *y*.

Based on the marginalization property of multivariate Gaussian distributions, the posterior covariance at a subset of points *z* can be obtained simply by taking the submatrix Σ_*zz*_ = *Z*^*T*^ Σ*Z*. We can verify that this also matches the classical solution for Gaussian process posterior covariance when *K* = *C*^−1^ using Woodbury’s identity:

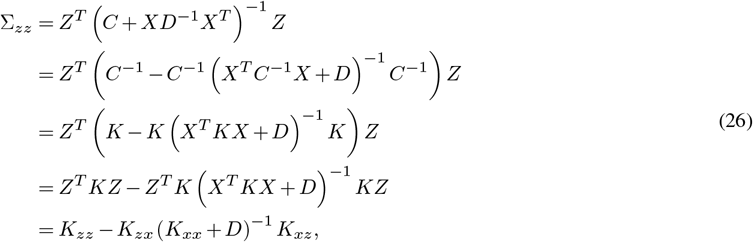

as desired.

